# Intracellular interactions shape antiviral resistance outcomes in poliovirus via eco-evolutionary feedback

**DOI:** 10.1101/2025.05.20.655113

**Authors:** Alexander J. Robertson, Benjamin Kerr, Alison F. Feder

## Abstract

Antiviral resistance evolution poses a major obstacle for controlling viral infections. A promising strategy is to target shared viral proteins that allow drug susceptible viruses to sensitize resistant ones during cellular coinfection, muting selection for resistance. Pocapavir, a poliovirus capsid inhibitor, employs this sociovirological strategy. While susceptible viruses significantly suppressed resistance in the presence of pocapavir in cell culture, a pocapavir clinical trial observed widespread resistance evolution and limited improvements to clearance times. To reconcile these findings, we present an intra-host eco-evolutionary model of poliovirus in the presence of pocapavir, which reproduces both the potent interference observed *in vitro* and the resistance emergence seen in patients. In the short term, our model predicts that a high density of susceptible viruses sensitizes resistant ones to pocapavir, mirroring cell culture results. However, over multiple replication cycles, pocapavir’s high potency collapses viral density, which reduces coinfection and allows resistance to evolve as observed in the clinical trial. Since coinfection is essential to suppress resistance, enabling greater survival of susceptible viruses could offer therapeutic advantages. Counterintuitively, we demonstrate that this can be achieved by *lessening* antiviral potency, which can limit resistance evolution while also maintaining a low viral load. These findings suggest that antivirals that rely on viral intracellular interaction must balance immediate neutralization with the preservation of future coinfection, yielding more sustained inhibition. Explicitly considering the eco-evolutionary feedback encompassing viral density, shared phenotypes and absolute fitness not only provides new insights into designing effective therapies but also illuminates viral evolutionary dynamics more broadly.

## Introduction

Resistance evolution can undermine otherwise successful antimicrobial treatments if drug application permits microbes carrying resistance alleles to expand and prevent population extinction. Implicit in these dynamics is that selection for phenotypic resistance also selects genotypic resistance, which permits the trait to carry forward into future generations. While such an assumption is often valid in bacterial populations (**Figure 1A-C**) [1, 2], in viruses, the association between genotype and phenotype can be more complicated [3, 4, 5]. For example, both cellular coinfection or *de novo* mutation during viral replication can result in distinct genotypes occupying the same cell. These genotypes may share intracellular protein products, and as a result, a particular genotype may be associated, fully or partially, with the protein products of a different genotype (**Figure 1D&E**) [6].

**Figure 1.**
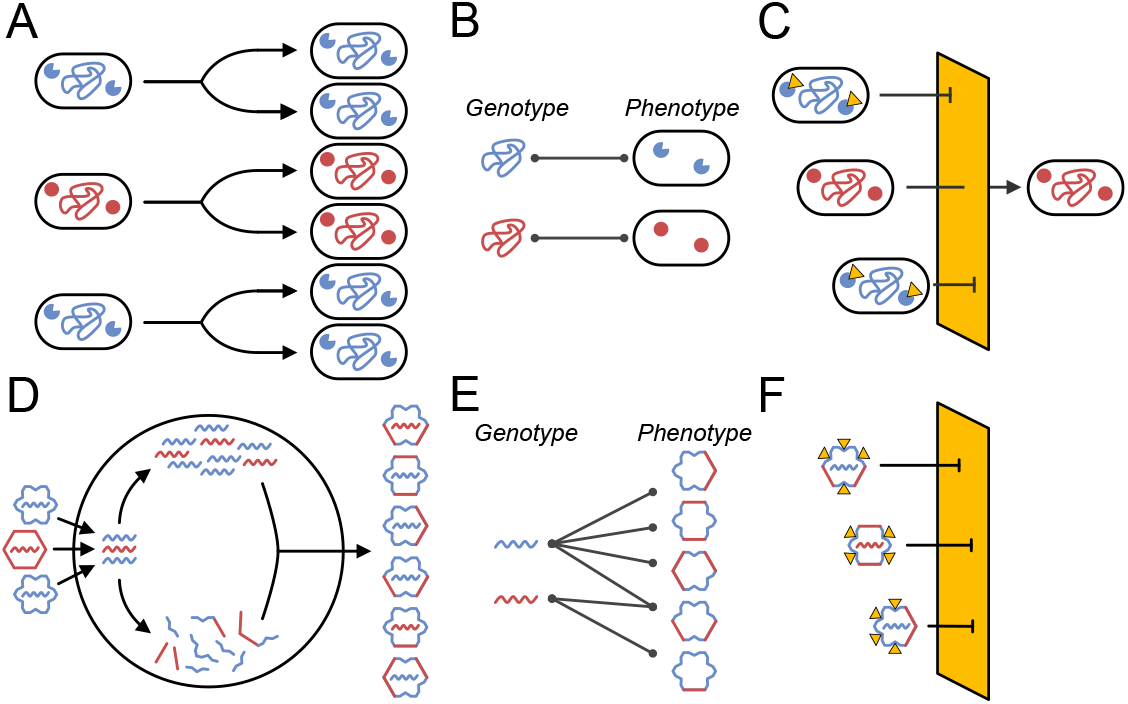
Genotype-phenotype associations can limit resistance evolution. In bacterial populations, resistance-conferring genotypes exist at low frequencies prior to treatment (**A**). Because genotypes are directly associated with the phenotypes they encode (**B**), drug application that selectively favors the survival of resistant phenotypes drives the expansion of resistant genotypes (**C**). In viruses, multiple genotypes (i.e., resistant and susceptible) may co-infect the same cell via mutation or superinfection and the protein products they encode can mix intracellularly (**D**). The resulting chimeric phenotypes can be associated with either genotype (**E**) and even those phenotypes that are partially composed of resistant proteins may be susceptible to drug neutralization (**F**) Throughout the figure, red and blue represent resistant and susceptible variants, respectively, and yellow triangles represent antimicrobials that can bind to susceptible proteins.

This so-called “phenotypic mixing” has important implications for resistance evolution [5, 7]. First, the disassociation between genotype and phenotype can impair selection for resistant genotypes. Consider a cell that contains both treatment-resistant and susceptible genotypes. If resistant genotypes do not associate with resistant protein products, those genotypes may not survive drug application, limiting their contribution to future generations (**Figure 1F**). Second, phenotypic mixing can limit the creation of resistant phenotypes themselves. If a drug targets an oligomeric protein product (e.g. a capsid inhibitor that targets the capsid), susceptible proteins can act in a dominant negative fashion to interfere with the phenotypic expression of resistance. Capsid inhibitors may neutralize chimeric capsids composed of both susceptible and resistant subunits if a sufficient number of susceptible subunits are available to bind the drug. If intracellular resistant genotypes are rare (such as after *de novo* mutation of a resistant mutant), cells may produce few or no phenotypically-resistant capsids. The presence of one or both of these factors could present an opportunity to treat viral infections while muting selection for resistance.

Exploiting phenotypic mixing to suppress resistance evolution while treating viral infections partially underlies the promise of the poliovirus capsid inhibitor pocapavir [6, 9]. Pocapavir strongly inhibits poliovirus, but mutations in the genes encoding capsid subunits VP1 or VP3 disrupt drug binding and confer resistance [10, 11]. When resistant viruses are co-cultured with susceptible ones and treated with pocapavir, resistant genomes are packaged in chimeric resistant-susceptible capsids and are neutralized by the drug [6]. As a result, co-cultured resistant viruses have significantly reduced viral titers compared to resistant viruses grown in isolation (**Figure 2A**). Promisingly, pocapavir was used to slow disease progression *in vivo* in poliovirus-infected mice with no detected resistance evolution [6]. However, in a larger placebo-matched human clinical trial where participants received the live attenuated polio vaccine and were treated with pocapavir, the drug both failed to significantly reduce time to viral clearance in three of four placebo-matched groups and resistance evolved in nearly half of pocapavir-treated participants (**Figure 2B**) [8]. While a subset of pocapavir recipients did clear their virus early without apparent resistance evolution (and pocapavir therapy has been successfully employed in certain compassionate use cases [12, 13]), results from the clinic have been mixed, and, to our knowledge, have not led to additional trials. More generally, this study raises doubts about the therapeutic potential of exploiting phenotypic mixing. To investigate these conflicting findings, we developed a dynamical model of poliovirus replication and evolution under drug treatment. Surprisingly, we find that a single model can reproduce the seemingly contradictory *in vitro* cell culture and clinical results via its behavior at different viral densities. At high viral density, susceptible viruses mask the phenotype of resistant ones and suppress selection for resistance, as observed in cell culture. However, as successful treatment drives the viral density down, limited intracellular viral interaction restores the standard genotypephenotype association. At this point, resistant and susceptible genomes associate strongly with their own phenotype, and drugs can efficiently select for genotypic resistance. Counterintuitively, this suggests that, in our model, permitting more susceptible viruses to survive drug application can better suppress resistance evolution and lead to smaller viral population sizes over time. This study provides the first theoretical framework for evaluating viral evolutionary responses to therapies targeting resistance phenotypes encoded by multiple genotypes, serves as a guide for the development of novel antimicrobials and dosing strategies, and highlights the importance of emergent dynamical responses when exploiting virus-virus interactions in medicine.

**Figure 2.**
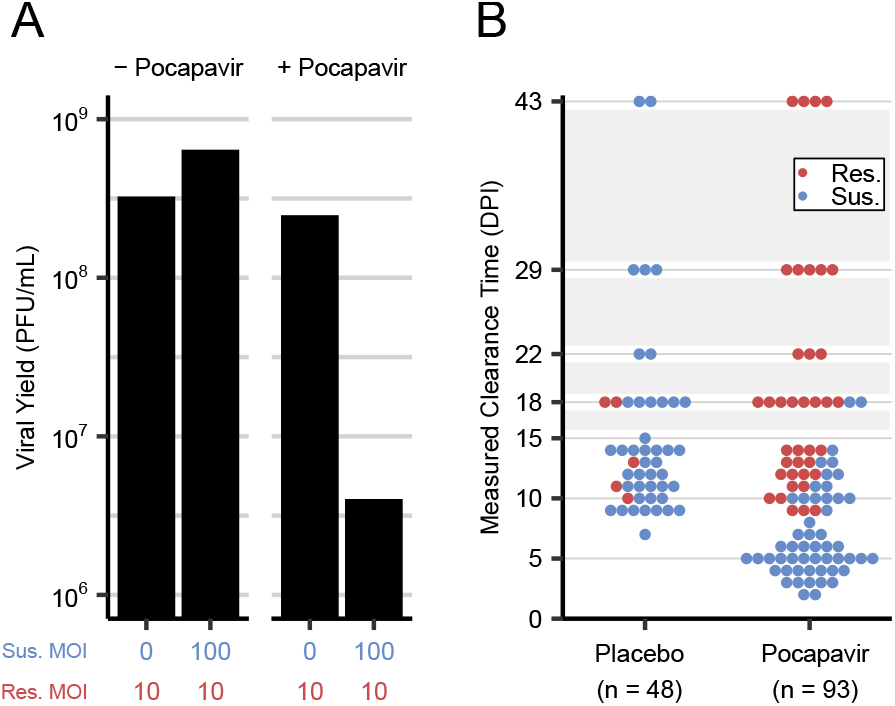
Pocapavir-treated poliovirus outcomes diverged between cell culture and clinical trial settings. (**A**) *In vitro* experiments by Tanner et al. [6] demonstrated that coinfection of drug-resistant (res.) and susceptible (sus.) poliovirus strains suppresses the yield of resistant virus under pocapavir treatment (MOI—multiplicity of infection, the ratio of viruses to host cells, PFU– plaque forming units). (**B**) In the clinical trial reported by Collett et al. [8], pocapavir failed to significantly reduce time to infection clearance compared to a placebo in three of four matched groups of participants administered the live attenuated poliovirus vaccine and resistance was enriched in the pocapavir group. Points represent the clearance dates of individual trial participants and are colored by resistance status (resistance in red, susceptible in blue) and gray boxes indicate dates that were not sampled during the trial (DPI—days post infection).

## Results

### Poliovirus eco-evolutionary model

We developed a discrete-generation dynamical model that tracks poliovirus genotypes and phenotypes over multiple rounds of viral replication and analyzed the model using both deterministic and fully stochastic simulations (see Materials & Methods). Briefly, each generation consists of four steps (**Figure 3A**):

**Figure 3.**
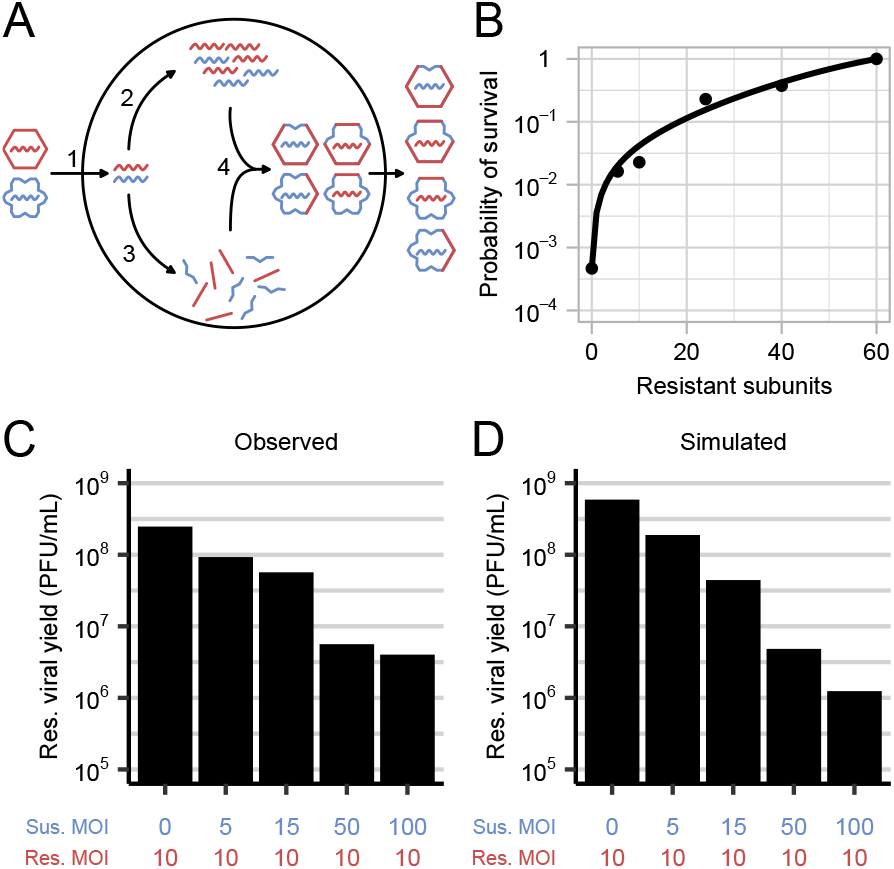
Discrete-time model of poliovirus replication, mutation and survival under pocapavir treatment. (**A**) We simulate intracellular poliovirus dynamics in four stages: (1) viral entry into host cells, (2) genome replication with mutation, (3) production of capsid subunits, and (4) assembly and packaging of progeny virions. (**B**) Capsid-mediated survival is modeled by culling progeny virions according to their capsid composition. Capsid survival probability as a function of the number of resistant subunits (line) was fit from cell culture experimental data from Tanner et al. [6] (points). Resistant viral yield under different intensities of susceptible virus coinfection shows a density-dependent effect *in vitro* [6] (**C**) and *in silico* under our model (**D**). Throughout the figure, resistant variants are illustrated in red and susceptible variants are illustrated in blue.

1. *Viral entry into host cells*: resistant and susceptible genomes enter cells as a function of their respective population sizes. Coinfection is more likely if viral population sizes are large relative to the host cell population.
2. *Viral genome replication and mutation*: intracellular viral genomes replicate up to the cell’s burst size. Mutation can interconvert resistant and susceptible genotypes.
3. *Capsid formation*: the initial infecting genomes produce a pool of shared capsid subunits. For mixed infections, capsids are formed by randomly drawing 60 subunits from this pool and can be composed of both resistant and susceptible subunits.
4. *Capsid packaging* : newly replicated viral genomes are packaged into assembled capsids in proportion to their intracellular abundances.

Viruses then exit cells and pocapavir can bind and neutralize free virions according to their capsid phenotype. We parametrized viral neutralization rates based on the reduction in viral titers measured experimentally in pocapavir-treated populations of mixed resistant and susceptible cultures (see Materials & Methods—Parameter inference). Capsids composed solely of resistant subunits survive pocapavir application with probability 1, while those composed of fully susceptible subunits survive with probability 4 × 10^−4^ (**Figure 3B**). Our model recapitulates the observations in Tanner et al. [6] that titers of resistant viruses (specifically, viral genomes encoding resistant subunits) decrease when co-infected alongside susceptible viruses (specifically, viral genomes encoding susceptible subunits) when treated with pocapavir (**Figure 3C&D**).

### Resistance suppression is dependent on susceptible virus density

We first assessed the conditions under which pocapavir resistance evolution is suppressed during a single round of replication with pocapavir treatment. Specifically, we measured the change in resistance frequency as a function of the total multiplicity of infection (MOI—a measure of viral density defined as the ratio of the total number of viruses to the total number of infectable host cells). We initialized simulations with the resistant genotype frequency set at f_Res_ = 10^−4^, consistent with levels observed in untreated poliovirus populations [6, 11].

At high MOIs (MOI ≈ 10^2^), genotypic resistance increased in frequency by less than 10^−3^ after a single round of replication in the presence of pocapavir (**Figure 4A**). These results are consistent with Tanner et al. [6] and the logic underlying phenotypic mixing. However, this resistance suppression did not extend to populations that were initialized at lower MOIs. At MOI ≤ 1, the resistant genotype frequency increased to more than 12% of the population in a single round of replication.

**Figure 4.**
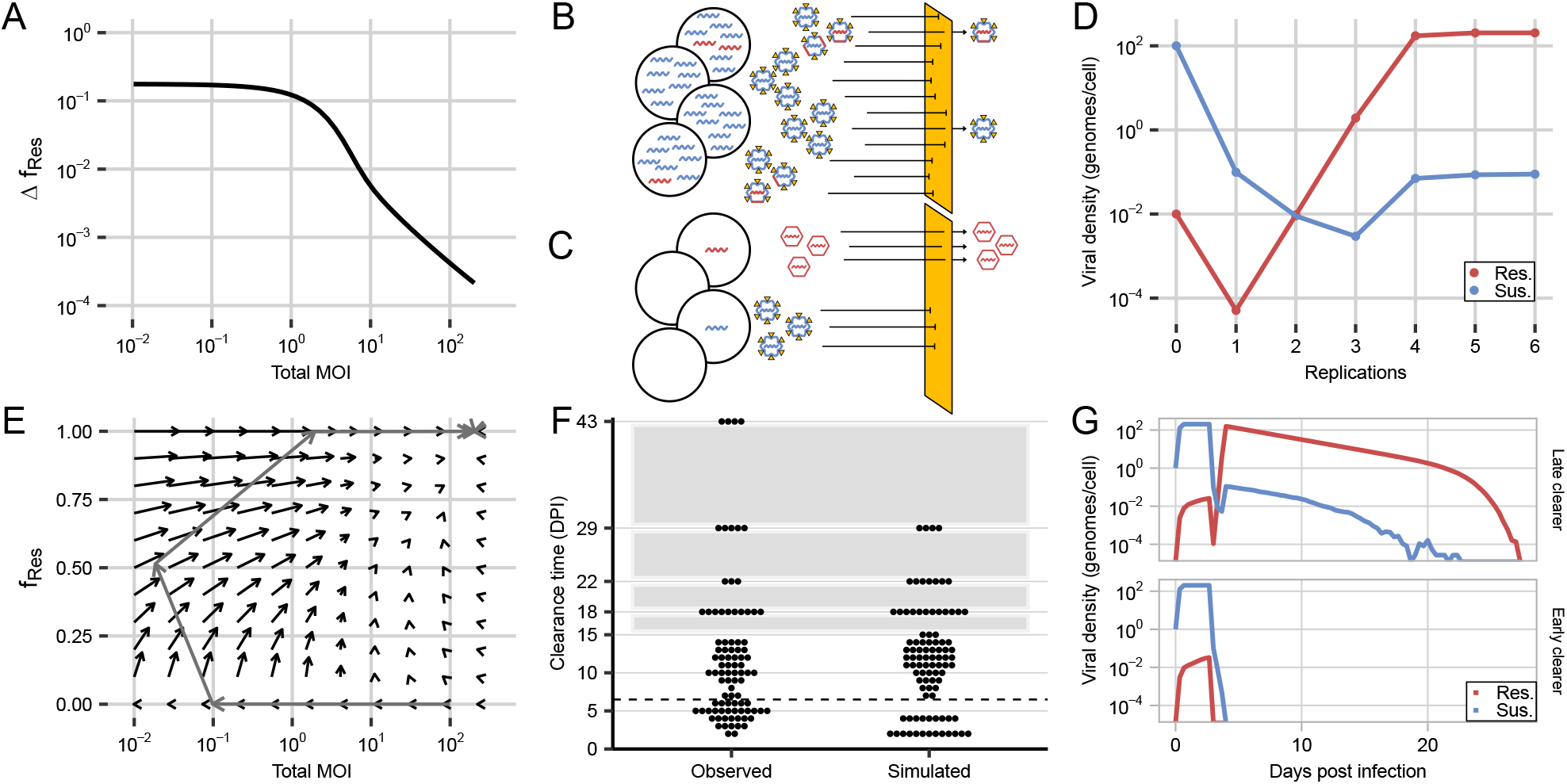
Resistance suppression is MOI-dependent and resistance emerges if bottlenecks do not lead to extinction. (**A**) Change in resistance frequency in a single generation (f_Res_) depends on the MOI (initial f_Res_ = 1 × 10^*-*4^). (**B**) At high MOI, rare resistant genomes are encapsidated by phenotypically susceptible capsids, muting genotypic selection. Although nearly all capsids are phenotypically susceptible, some survive pocapavir administration (a fraction greatly exaggerated for this cartoon), as observed in cell culture. (**C**) At low MOI, viruses singly infect cells, and rare resistant genomes are encapsidated by phenotypically resistant capsids, enabling selection for resistance. (**D**) Over multiple generations, pocapavir treatment transiently reduces viral population size for both resistant and susceptible genomes but leads to viral rebound of a primarily genetically resistant population following low population density. (**E**) Discrete step phase diagram shows the joint change in genotypic resistance frequency and MOI from different initial conditions. Arrows are shortened by 80% to increase legibility. The trajectory from part (**D**) is overlaid in grey. (**F**) Clearance dates from the observed clinical trial and one simulated clinical trial of n = 93 viral populations treated with pocapavir. The dashed line indicates the date of the earliest placebo clearance. (**G**) Late and early clearers both experienced drops in resistant and susceptible viral population size with diverging outcomes following the population bottleneck. Throughout the figure, resistant variants are illustrated in red and susceptible variants are illustrated in blue.

These results can be understood in the context of the MOI controlling the degree of coinfection, and subsequently the strength of the genotype-phenotype association. At high MOIs, coinfection is ubiquitous and the association between phenotypes and rare genotypes breaks down, and rare resistant genomes are wrapped in susceptible capsids (**Figures 4B, S1**). However, at low MOIs, coinfection is rare and genotypes and phenotypes are directly associated (**Figure 4C**), allowing resistant genotypes to directly benefit from their associated phenotype without interference.

We next considered that infections are dynamic processes, and the degree of viral suppression or proliferation in one generation may determine viral density in subsequent generations. If pocapavir treatment drastically reduces the viral density (and consequently the total MOI), this may lead to conditions in which resistance can emerge. We therefore performed an *in silico* serial passaging experiment in which we inoculated cell populations at a high MOI (MOI = 100) with predominantly susceptible viruses (resistant genotype frequency f_Res_ = 1 × 10^−4^). We allowed viruses to replicate and be neutralized by pocapavir, and seeded the surviving viruses on fresh cell populations over multiple generations.

Consistent with single step experiments at high MOI, the viral population decreased in abundance after one round of replication in the presence of pocapavir (**Figure 4D**). As a result, the surviving viral progeny infected cells at significantly reduced MOI and resistant genomes increased in both frequency and abundance after a second and third round of passaging. At the passage at which the viral population size recovered enough for widespread coinfection, genotypic resistance was more than 100 × more common than genotypic susceptibility and susceptible subunits no longer effectively sensitized their resistant counterparts (**Figure S2**). Rather, resistant viruses appeared to shield rare susceptible genomes in a form of socially-encoded cross protection. These dynamics can be understood by tracking the change in MOI and resistance frequency on a stepwise phase diagram, initiating simulations across a range of initial total MOIs and resistance frequencies (**Figure 4E**). Initial conditions with low resistance frequencies and sufficiently high MOI lead first to a rapid reduction in the MOI, which then permits increases in the genotypic frequency of resistance. Note, one way to understand this sequence is that the low viral titer output of **Figure 4B** is the low MOI input of **Figure 4C**, enabling the spread of resistant viruses. Regardless of the initial conditions in our deterministic model (MOI > 0, f_Res_ ∈ [0, 1]), resistance becomes the dominant genotype over time. Despite phenotypic mixing effectively suppressing resistance at high MOIs, rapid population contraction in response to pocapavir ultimately undermines that suppression.

### Stochastic model replicates clinical trial outcomes

While our deterministic model can explain how resistance to pocapavir may have emerged in clinical trial participants despite phenotypic mixing, it can not recapture the clinical trial observation that a subset of pocapavir recipients clear their infections early with little to no resistance evolution (**Figure 2B**) [8]. To investigate the dynamics driving the clinical trial clearance times, we implemented a stochastic version of the model in which simulated viral elements are drawn from probability distributions at each replication step in a finite host cell population with an immune system that responds to infection (see Materials & Methods—Immune clearance). In brief, we modeled viral immune clearance via a non-specific, ramping innate immune response that removes viruses irrespective of capsid phenotype, and parameterized clearance rate and host cell population size based on the clearance dates in the clinical trial placebo group. We used the stochastic immune model to simulate an *in silico* pocapavir clinical trial by running 93 simulations (representing 93 trial participants) until viral extinction. The infections were initialized with one susceptible virus per host cell, and no resistant viruses. Pocapavir was administered after 24 hours (three rounds of replication, n = 23) or 72 hours (nine rounds of replication, n = 70), as in the clinical trial.

Our model broadly recaptures the clearance time and resistance evolution outcomes observed in Collett et al. [8]. Specifically, viral populations exhibit a bifurcation of outcomes, in which they clear shortly after pocapavir initiation (< 7 days post infection) with little genotypic resistance, or clear later (≥ 7 days post infection) with widespread genotypic resistance (**Figure 4F**). Analysis of the simulated population trajectories revealed that both early clearers and late clearers experienced sharp population bottlenecks and low MOIs shortly after pocapavir initiation (**Figures 4G, S3**). In late clearers, this bottleneck allowed for rapid, resistance emergence according to the dynamics explored above, while in early clearers, this bottleneck led to stochastic extinction. Repeating clinical trials with different host cell population sizes led to changes in the relative probabilities of these two outcomes (**Figure S4**), but not their qualitative behavior.

Collett et al. [8] also observed that a greater proportion of participants treated at 24 hours exhibited resistance than the 72 hour treatment group (15/23 versus 25/70, Fisher’s exact test, *p* = 0.0163), which is potentially unexpected given that more resistant genomes are expected to exist after 72 hours. Under certain starting conditions in our model, this pattern can emerge as resistant genomes produced before ubiquitous coinfection can be more tightly linked to their resistant phenotype counterbalancing their lesser numbers (**Figure S5**). In sum, our model can explain counterintuitive and divergent participant outcomes among pocapavir recipients in Collett et al. [8].

### Susceptible dominance has a limited effect on resistance suppression

We hypothesized that pocapavir resistance may evolve in poliovirus due in part to the particular relationship between the number of resistant subunits in a capsid and the virion’s fitness. Specifically, when we parameterized the pocapavir neutralization curve using experimental measurements from Tanner et al. [6], we found that the largest relative increases in capsid survival probability accompanied the incorporation of the first few resistant subunits (**Figure 3B**). We hypothesized that resistance could be successfully suppressed over longer timescales if more resistant subunits were required to increase the probability of a capsid surviving pocapavir (i.e., the susceptible subunits exerted a greater effect over phenotype than resistant subunits). We therefore repeated our experiments using a neutralization curve in which a virus needed to construct a fully resistant capsid (i.e., 60/60 resistant subunits) before it experienced any survival improvement in the presence of the drug (**Figure S6A**).

Intriguingly, under these conditions, dynamical outcomes were nearly identical to those under the neutralization curve of pocapavir, despite the seemingly more favorable properties. Again, resistance suppression was highly dependent on MOI (**Figure S6B**) and resistance evolved over multiple passages following a drug-induced reduction in viral population size (**Figure S6C**). More broadly, we found that as long as the survival probabilities of fully resistant and fully susceptible capsids were preserved (i.e. the end points of the fitness curve remained the same), a range of monotonically increasing functions describing the dominance interactions between susceptible and resistant subunits resulted in very similar clinical outcomes to pocapavir (**Figure S6D-F**). This consistent result emerges because the rapid contraction of the population following drug application results in most cells being infected by single viral genomes. At that point, interaction properties between resistant and susceptible subunits in the presence of the drug become secondary to the survival probabilities of fully resistant and susceptible virions. This suggests that efforts to find drug formulations in which resistance mutations behave more “recessively” are unlikely to yield substantively different results as long as drug application continues to produce singly-infected cells.

Potentially more impactful in outcome would be a hypothetical fitness cost to resistance. We find that simulating such a cost in our model slows resistance evolution, decreases the proportion of resistance at equilibrium, and causes excess stochastic clearance during population bottlenecks (**Figure S7**). However, this effect is unlikely to be clinically relevant here because pocapavir resistance mutations do not appear to be costly to poliovirus in cell culture [6, 10, 11].

### Reduced drug potency can enhance long-term control of resistance and lower viral burden

The critical liability of pocapavir identified above is that effective neutralization of the virus disrupts susceptible viruses’ abilities to interfere with resistant phenotypes via coinfection. We hypothesized that increased survival of susceptible viruses would enhance resistance suppression by maintaining higher rates of coinfection over time. Furthermore, since much of the viral burden in a resistance-dominated infection is driven by high viral loads following the emergence of resistance, effectively suppressing resistance could reduce the viral load overall, even if more susceptible viruses survive. We therefore considered the effects of hypothetically less potent drugs (or alternatively, lower doses of pocapavir) that permitted greater degrees of survival by virions with susceptible or partially susceptible capsids (**Figure 5A**, see Materials & Methods—Variation in drug strength).

**Figure 5.**
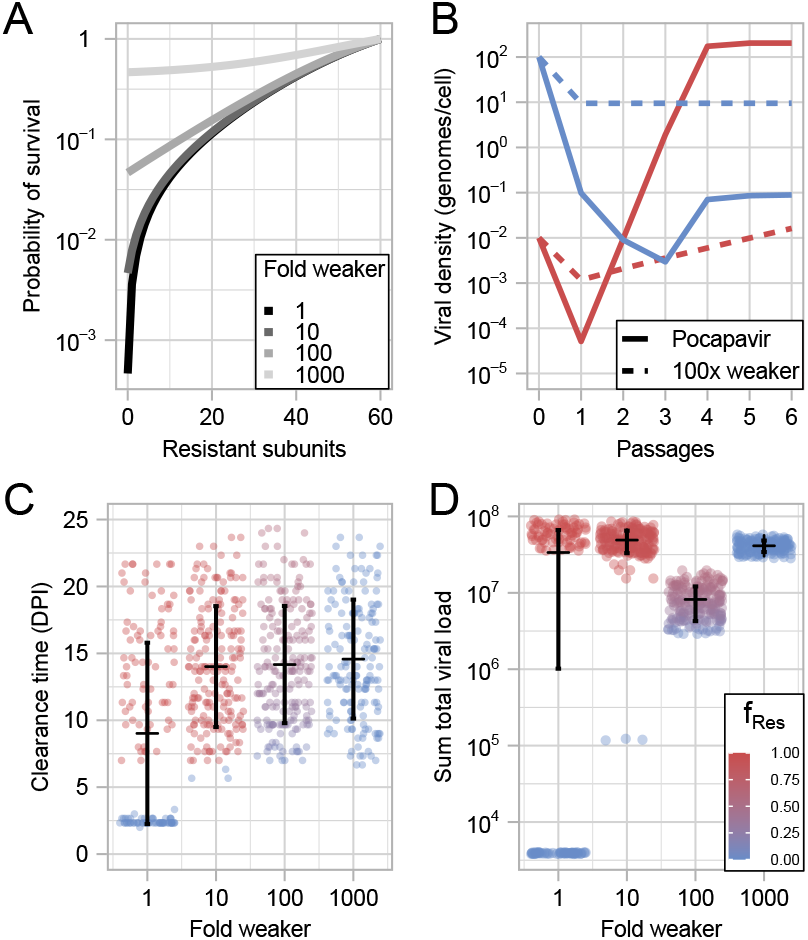
Drugs less potent than pocapavir can better suppress resistance and maintain lower viral loads. (**A**) We modeled hypothetical drugs 10×, 100× and 1,000× weaker than pocapavir and plot the probability of virion survival as a function of the number of resistant capsid subunits under these drug conditions. (**B**) In deterministic serial passage experiments, the 100× weaker drug maintained a lower total viral population than pocapavir (resistant genomes, red; susceptible genomes, blue). We simulated clinical trials of 100 individuals treated with drugs 1×, 10×, 100×, and 1,000× weaker than pocapavir. Reducing drug potency delayed mean clearance time measured by days post infection (**C**) but had a non-monotonic effect on the sum viral load over the course of the infection (**D**). In both (**C**) and (**D**), dots represent individual simulations and are colored based on the frequency of resistance in the population over the course of the infection (scale in (**D**)). Error bars show mean (red triangle) *±* variance. For an expanded range of drug potencies, see **Figure S8 B&C**.

Reducing drug potency led to smaller increases in the frequency of genotypic resistance in a single round of replication than pocapavir, in line with evolutionary expectations for weaker selective pressures (**Figure S8A**). However, reducing drug efficacy also led to a less rapid decline in the absolute number of susceptible genomes. For example, a drug with a 100-fold reduced efficacy reduced the MOI after a single round of replication to approximately 10, whereas pocapavir reduced the MOI to substantially less than one virus per cell (**Figure 5B**). As a result of this sustained moderate MOI, genotypic resistance increased minimally (up to 0.16% of the population) over the next 6 passages while genotypic resistance under pocapavir reached near 100%. Because resistance remained suppressed, the population did not undergo full viral rebound and the total MOI after 6 passages was approximately 20 times smaller under the 100× less potent drug than pocapavir. This suggests that reducing antiviral potency can, under some circumstances, improve multiple aspects of viral control.

To evaluate the potential clinical implications of reducing antiviral potency, we simulated *in silico* clinical trials of 100 individuals treated with reduced potency drugs (**Figure 5C&D**, assessed across a greater range in **Figure S8B&C**). We found that pocapavir had an earlier mean clearance time compared to the less potent hypothetical drugs. Reduced drug potency weakened population bottlenecks, preventing stochastic extinction of viral populations at small population sizes when treated with drugs weaker than pocapavir.

In contrast, reducing drug potency had more complex effects on the frequency of resistance and the sum total viral population size over the course of infection (**Figure 5D**). Reducing drug potency by a factor of 10 resulted in near ubiquitous resistance evolution and high viral population size across trial participants. This reflects that a 10× reduced potency is a strong enough selective pressure to bring viral populations into a regime in which cells are singly infected and resistance can evolve, yet it is not strong enough to create population bottlenecks severe enough for stochastic extinction like pocapavir can. In contrast, reducing drug potency by a factor of 100× created a gradual enough decay during initial drug application that resistant-susceptible coinfection was durably maintained. While this rate of viral decay was not strong enough to cause stochastic extinction during bottlenecks, the sustained coinfection reduced the rate of resistance evolution and subsequently the total viral load compared to pocapavir. Finally, a very strong reduction of drug potency by a factor of 1, 000× selected for almost no resistance evolution, but also did not

restrict the viral population relative to an untreated control. This suggests that while reducing drug potency can improve multiple important clinical metrics within the context of our model, not all reductions in drug potency will have favorable effects and there is an optimal balance between preserving susceptible virus and limiting total infection burden.

## Discussion

In this paper, we show that a single model of poliovirus population dynamics and genetics can reconcile seemingly divergent outcomes of pocapavir treatment in cell culture [6] and clinical trial settings [8]. Our key insight is that therapeutic strategies that rely on interaction between viruses must account for the demographic effects of therapeutic success. If the long-term efficacy of these therapies depends on the durability of intracellular interactions, lowering viral density through successful treatment can decrease coinfection rates and thus the potential for therapeutically beneficial interactions between viral genomes and their encoded proteins.

These considerations are important given the growing interest in exploiting intracellular interactions between viral genomes in a therapeutic capacity. In most of these proposed cases, partially or fully-defective viral genomes serve as the therapeutic themselves rather than a sensitizing agent for a different small molecule as we consider here. Perhaps most promisingly, therapeutic interfering particles (TIPs) are replication-incompetent viral mutants that can interfere with their replication-competent counterparts when they co-infect the same cells [14, 15, 16]. Because TIPs expand faster than their targets upon coinfection, their interference capability grows rather than shrinks over time (as long as conditions favoring coinfection endure). Although the full therapeutic potential for TIPs is still an active area of investigation, Pitchai et al. [17] recently demonstrated an effective proof of concept for using TIPs to treat HIV over short timescales. The demographic feedback considered in our model highlights potential challenges with these approaches - over longer periods of time, if TIPs drive replication-competent viruses to near but not complete eradication, TIPs may lose their ability to self-renew and themselves become eliminated. This could ultimately lead to the unencumbered rebound of replication-competent viruses, especially in the case of viruses with a latent stage such as HIV where viruses can reactivate from a reservoir after TIPs have been eliminated.

If intracellular interactions are crucial for the success of these treatments, how can they be maintained in the face of demographic collapse? In the case of TIPs, Weinberger, Schaffer, and Arkin [15] found that *weaker* interference between TIPs with their wild-type counterparts favors the long-term success of TIP therapy [15]. Indeed, all successful demonstrations of TIPs in rodents [18, 19, 20] or mosquitoes [20] allow sufficient wild-type replication to maintain therapeutic pressure in subsequent generations. In our model of poliovirus and pocapavir, we achieve sustained intracellular interaction through a similar therapeutic intervention—decreasing drug intensity. Modifying drug intensity through dosage or administration frequency may allow more fine-tuned calibration of intracellular interactions than identifying TIPs with the appropriate degree of interference. To be clear, this is not a clinical recommendation, and many important model assumptions would need to be rigorously evaluated in experimental settings before altering any therapeutic approach. That being said, similar strategies to prolong drug efficacy by exploiting interactions between different genotypes are also being explored in non-virology settings. For example, in drug-treated bacterial [2, 1] and cancer [21, 22] populations, moderate-dose or pulsed strategies can prolong competition between susceptible and resistant cells, thereby lowering overall population size over time and delaying treatment failure.

Although these treatment strategies reckon with similar demographic dynamics across systems, they may require different medical and public health considerations. In our model, decreased drug potency can slow the emergence of resistance and lower viral load. However it simultaneously prolongs average infection clearance time by reducing stochastic early viral extinction. Similarly, in cancer adaptive therapy, alternating therapeutic pressure and drug holidays reduces tumor burden and increases the time to clinical progression, but sacrifices the possibility of eradication and ultimately only delays resistance evolution [21]. While both treatment strategies therefore affect individual patient clinical outcomes, infectious diseases carry the added risk of onward transmission. Decreased viral load from partial population control may result in reduced poliovirus infectivity as it does in SARS-CoV-2 [23, 24] and HIV [25], but slower clearance times may counterbalance this effect by increasing the number of contacts exposed. Resistance evolution also becomes a more acute concern than in cancer therapies, as resistant variants can transmit to other people and undermine future drug effectiveness. To determine if the decreased viral load and enhanced resistance suppression of a less potent drug outweighs the concomitant increase in the infectious period, an epidemiological model incorporating the intra-host dynamics we explore could prove valuable.

While intracellular interaction is a liability to viruses in our model, these same interactions may be beneficial or even critical to viruses in other settings. For example, multipartite viruses package their genome segments into separate capsids and therefore require cellular coinfection of virions carrying distinct segments to successfully replicate [26]. Coinfection may also be critical for influenza viruses, as individual virions often carry incomplete genomes [27, 28]. In less extreme cases, increased intracellular interactions can allow viral populations to more effectively explore fitness landscapes [4, 3, 7] and exchange genes [29, 30]. When considering viral evolution in these settings, positive demographic feedback loops may need to be considered.

A central assumption in our model is that the number of viruses per host cell is the primary driver of coinfection rate. While this is a common assumption [31, 32, 33, 4], diverse viral behaviors can modulate coinfection. For example, the first virus infecting a cell can prevent others from entering in a process called superinfection exclusion [34]. Conversely, *en bloc* transmission, in which viruses are packaged and transmitted collectively, can enhance coinfection [35, 36]. To our knowledge, poliovirus does not have superinfection exclusion behavior [37, 38], but growing evidence suggests that *en bloc* transmission [35, 39, 36] and even shuttling of poliovirus by enteric bacteria [40] can be common during infection. These factors could elevate poliovirus coinfection rates beyond what we consider in our model, or change which genotypes coinfect together.

Host and environmental factors could also affect the realized frequency of coinfection and subsequent evolutionary dynamics. We assume that cells are all equally susceptible to poliovirus infection and spatially well-mixed. In practice, expression of poliovirus’ primary receptor CD155 varies considerably among cells [41] and at different stages of disease [42]. This could concentrate virus into a smaller number of cells, enhancing interference, or result in certain cells that can only be infected by few virions, potentially allowing greater expression of phenotypic resistance. The organization of host cells within a tissue also stands to impact coinfection dynamics [43]. Limited viral dispersal could increase coinfection and therefore interference, but it might also concentrate resistant genomes into the tissue sections in which they initially arose, limiting the degree to which susceptible genomes could interfere with resistant spread. The degree of this effect might also vary across organ systems. Notably, the limited resistance evolution observed in pocapavir-treated mice may relate to increased viral density in neuronal infections [6, 44] relative to the gut epithelial infections of the pocapavir clinical trial in which resistance evolution was common. Investigating the importance of intra-host spatial and environmental variation is an important future area of research.

A second potentially important form of spatial organization is intracellular. Many viruses, including poliovirus, form membrane-associated structures that can sequester viral components near their encoding genomes [45, 46]. This effect could limit protein diffusion and reduce phenotypic mixing. Poliovirus capsids are tethered to their membranes during virion assembly [47], so it is possible that some degree of intracellular segregation contributes to our observation that even very rare resistant genomes in a cell impart partial phenotypic resistance. Despite this, Tanner et al. [6] observed that phenotypically-distinct capsid proteins intermingle in single chimeric virions, so the exact extent of intracellular mixing of poliovirus capsid subunits remains unknown. Nevertheless, the organization of viruses within a cell is likely to be a key determinant of therapeutic strategies built around intracellular resource sharing.

More broadly, our study contributes to a growing body of work framing viral coinfection through the lens of ploidy and drawing parallels to how multiple gene copies shape phenotype in cellular organisms [48, 7, 49, 50]. In classical diploid genetics, a single biallelic locus can produce up to three phenotypes, depending on the interactions between the alleles (i.e., the “dominance” of one allele over the other). Although the frequency of these phenotypes can shift between generations, the ploidy level itself remains fixed. Among viruses, the number of genomes that contribute to a phenotype can vary between host cells and dynamically over the course of an infection [7]. The association of several viral genomes in a host cell can lead to more complex and varied phenotypes than is possible in a diploid model [51]. The model that we explore here has a classical analog to incomplete dominance in a standard diploid framework, in which the addition of more resistant proteins always partially benefits a capsid in the presence of the drug. However, recent work has also described instances of apparent over- or underdominance in which coinfecting viruses have increased or decreased fitness relative to either non-mixed infection [52, 53, 54, 55]. Regardless of the exact form of intracellular “dominance,” shared phenotypes can clearly determine viral fitness. Because absolute fitness within a population governs viral density in the immediate future, it therefore impacts the degree of viral intracellular interaction moving forward, and thus how new phenotypes are realized. Therefore, viral “ploidy” not only changes over time, but actually feeds back into itself (**Figure 6**). Although our model centers on poliovirus and pocapavir, its core principles—density-driven coinfection, transient genotype-phenotype associations, and demographic feedback with the environment —are likely to be broadly relevant for designing therapies that exploit the social lives of viruses, and viral evolutionary dynamics more broadly.

**Figure 6.**
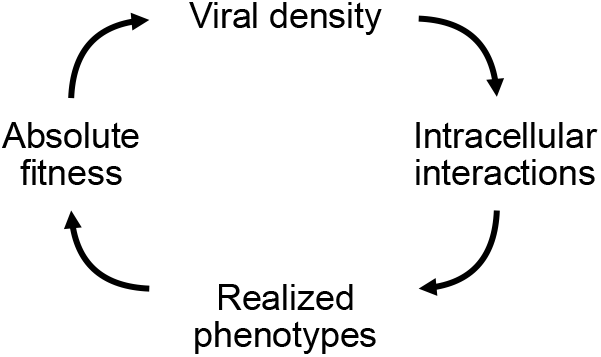
Targeting viral traits affected by multiple intracellular viral genomes requires an understanding of eco-evolutionary feedback. Viral density determines the degree of intracellular interactions, which determines realized phenotypes, which determine absolute fitness, which feed back into viral density in the next generation.

## Materials & Methods

We developed both stochastic and deterministic versions of a discrete-time dynamical model of poliovirus replication that integrates viral entry, genome replication, mutation, capsid formation, pocapavir neutralization, and immune clearance. We describe the stochastic version here, but the deterministic version follows the same probability distributions but accounts for each possible outcome (i.e., integrates over the probability distribution) instead of drawing specific values. Full details are provided in the Supplemental Materials & Methods. All simulations and analyses were performed in R, version 4.1.0 [56]. Data visualization was performed with the ggplot2 package [57].

### Viral entry

At time *t*, the total viral population, *v*_tot,*t*_, consists of *r*_tot,*t*_ resistant and *s*_tot,*t*_ susceptible genomes, yielding

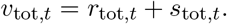

We assume that viruses are equally likely to infect any of the γ host cells in the population, and that there is no superinfection exclusion. The number of resistant and susceptible genomes entering a cell (random variables *R*_inf_ and *S*_inf_, respectively) are modeled by binomial distributions,

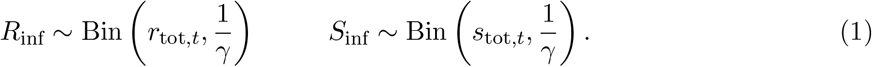

The total number of viruses that have infected a given cell, *v*_inf_, can be described by the equation: *v*_inf_ = *r*_inf_ + s_inf_, where *r*_inf_ and *s*_inf_ represent realizations of the binomial distributions described in Equation 1. Thus, on average, *v*_inf_ will equal 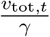, the multiplicity of infection (MOI).

We note that in the deterministic model, *r*_tot,*t*_ and *s*_tot,*t*_ need not be integers, rendering the binomial distribution undefined. We describe a weighted sampling scheme to circumvent this issue in Supplemental Materials – Viral Entry.

### Genome replication and mutation

For each infected cell, progeny production (*V*_rep_) follows a Poisson distribution with a mean of the inferred average effective burst size, *β*, which accounts for the number of infectious particles that leave the cell (i.e., *V*_rep_ ∼ Pois(*β*) for *v*_inf_ > 0). Replicated resistant genomes (*R*_rep_) are modeled as

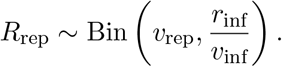

Given that *R*_rep_ takes on some value, *r*_rep_, the number of newly replicated susceptible genomes in a cell, *s*_rep_, is given by:

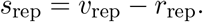

Mutation between genotypes occurs per replication event at rate *µ* = 2 × 10^*−*5^ [10], so that resistant and susceptible mutants (the random variables *R*_mut_ and *S*_mut_, respectively) are found by:

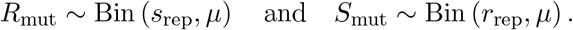

Given that *R*_mut_ and *S*_mut_ take on the values *r*_mut_ and *s*_mut_, respectively, post-mutation genome counts per cell (*r*_pool_ and *s*_pool_) are then found as follows:

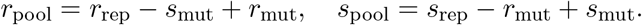

### Capsid formation

Let *σ* represent the number of subunits in a capsid. Each progeny genome is packaged into a capsid comprised of 60 subunits (*σ* = 60). The number of resistant subunits per virion (the random variable *I*) is modeled binomially, where:

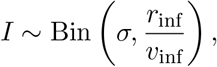

assuming that both resistant and susceptible genomes contribute equally to the pool of capsid subunits. The probability that a capsid has *i* subunits is:

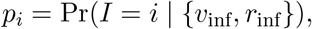

where *i* takes on a discrete value between 0 and *σ*.

Genomes are assigned to capsids via a multinomial sampling process for each infected cell in the population. The number of replicated resistant genomes that are packaged into a capsid with i resistant subunits is the random variable *R*_pack,*i*_. Each *R*_pack,*i*_ can be collected into a vector *R*_pack_, where:

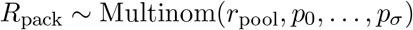

with analogous sampling for susceptible genomes. At this point, virions leave their cells and are pooled into groups according to their capsid subunit composition and genotype.

### Pocapavir neutralization

Drug neutralization is modeled by assigning each virion a survival probability, ω(*i, t*), that depends on its capsid composition (number of resistant subunits i) and on the time of drug administration *t*_poc_. For t < *t*_poc_, ω(*i, t*) = 1 (that is, before drug application, virions are not affected by the drug). When *t* ≥ *t*_poc_, survival is given by a scaled logistic function:

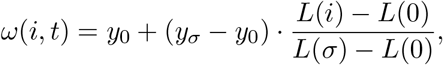

where *L*(*i*) is the standard form of the logistic function:

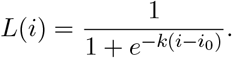

The variables *y*_0_ and *y*_*σ*_ represent the survival probabilities of fully susceptible and fully resistant capsids, respectively, and *k* and *i*_0_ are inferred by fitting the function to cell culture survival probabilities from Tanner et al. [6]. Under drug pressure, the survival of virions carrying resistant genomes (the random variable *R*_surv,*i*_) is then found by binomial sampling *r*_pack,*i*_:

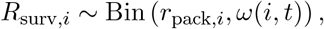

for each capsid subunit state (and similarly for susceptible virions). *R*_surv,*i*_ then takes on the specific values *r*_surv,*i*_. After pocapavir neutralization, the total number of resistant genomes, *r*_sum_, is calculated by summing across all *r*_surv,*i*_:

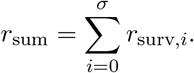

A similar sum is used to calculate total number of susceptible genomes.

### Immune clearance

Following pocapavir neutralization, immune clearance is applied after a specified number of replications, *t*_imm_, via the survival function d(t). For *t* < *t*_imm_, *d*(*t*) = 1. For *t* ≥ *t*_imm_, survival is given by an exponential decay function,

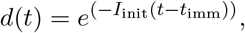

where the initial immune sensitivity, *I*_init_, is drawn from a lognormal distribution,

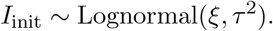

The parameters governing *I*_init_ (*t*_imm_, ξ and τ) and the host cell population size, γ, were inferred by maximizing the log-likelihood of observing simulation clearance times, given matched placebo clearance times from the pocapavir clinical trial reported by Collett et al. [8].

The virions that survive pocapavir treatment and carry resistant genomes are represented by *R*_imm_. This is found by binomially sampling *r*_sum_:

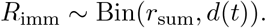

Note, immune clearance is considered in the stochastic model only.

### Initializing the next generation

In the stochastic model, draws from *R*_imm_ and *S*_imm_ represent the number of resistant and susceptible genomes that can infect cells in the next generation (i.e., *r*_tot,*t*+1_ and *s*_tot,*t*+1_). If the realized values of r_tot,*t*+1_ and s_tot,*t*+1_ both equal zero, the simulation is terminated.

### Variation in phenotypic dominance

To explore the impact of susceptible dominance over the resistant phenotype, we changed the drug neutralization function to simulate different relationships between resistant subunit composition and virion survival. Using the standard logistic function as our base, we used a steepness coefficient of *k* = 100 to simulate a step-like function, and varied *i*_0_ to set the inflection point, corresponding to the minimum number of resistant subunits needed to render a virion phenotype resistant to drug. Otherwise, simulations were initialized with the same parameters as the pocapavir simulations.

### Fitness cost of resistance

We examined fitness costs of resistance through a linear fitness function in which each additional resistant subunit in a capsid was associated with a κ decrease in virion extracellular survival probability, regardless of drug pressure (κ ∈ [0, 0.0165], **Figure S7**). Unless otherwise noted, κ = 0.

### Variation in drug strength

To explore the impact of drug strength on clinical outcomes, we scaled the original pocapavir fitness function such that,

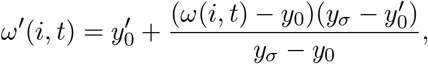

where *y*_0_ is the fitness of a fully susceptible capsid in the presence of pocapavir, *y*_*σ*_ is the fitness of a fully resistant capsid in the presence of pocapavir, and 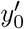 is the new survival probability of a virion composed entirely of susceptible subunits. Otherwise, simulations were initialized with the same parameters as the pocapavir simulations.

### Parameter inference

Parameters were inferred by numerical optimization in R (version 4.1.0) [56]. The specific objective function varied by model and is described below.

### Estimation of viral burst size

We inferred the effective viral burst size by fitting model outputs to mixed cell culture data from Tanner et al. [6], using matched initial multiplicities of infection (MOIs). We specifically compared our simulated data to reported results of pure Mahoney strain PV with the VP3-A24V mutation in cell culture. This mutation is one of the most commonly observed in experiments selecting for resistance [10, 11], and was the only strain/mutation pairing for which there was a negative control reported in Tanner et al. [6]. We evaluated model fit by minimizing the sum of squared differences between the log-values of the observed and simulated PFU after one round of replication. Specifically, we minimized:

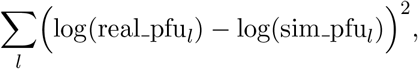

where *l* is the set of resistant MOI and susceptible MOI co-infected pairs reported by Tanner et al. [6]. Fit values and their empirically measured comparisons are reported in Table S1.

### Estimation of immune clearance parameters

Parameters governing immune clearance and host cell population size were estimated by fitting a stochastic model of infection and clearance (assuming no drug effect; i.e., survival = 1 for all phenotypes at all time points) to placebo group clearance data from Collett et al. [8]. Since participants were not sampled daily, Collett et al. reported the clearance date as the first sample at which no virus was detected (**Figure 2B**). To replicate this, we rounded each simulated clearance time up to the next available sampling day, following the trial design.

We simulated 480 placebo recipients (10× the original sample size) and calculated the probability of clearance on each sampled day. These model-based probabilities were compared to the empirical distribution using a multinomial log-likelihood. Specifically, after adding a small pseudocount to avoid log-zero issues, we computed:

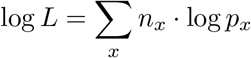

where n_*x*_ is the number of placebo participants observed to clear on day *x*, and *p*_*x*_ is the modelderived probability of clearance on day *x*. If a simulated participant cleared after day 43 (the final sampling day), we could not compare this outcome to empirical data and instead returned a fixed log-likelihood value of 1000 to penalize these parameter settings. Fit values and their empirically measured comparisons are reported in Table S1.

We note that the relevant *in vivo* cellular population size γ is not well-described in the literature, and its fit value in our optimization routine is sensitive to starting conditions, suggesting that it does not drive likelihoods. We present results with the fit value of *γ* = 37, 041 in the main text, as it broadly matches clearance outcomes in the treated group, with two additional *γ* values resulting from different optimization initial conditions in the Supplement (**Figure S4**). We further analytically characterize the dependence of the extinction probability on *γ* × *µ* in Supplemental Materials & Methods – *In vivo* cellular population size and **Figure S9**.

A complete account of the model equations, parameter inference, and simulation details is provided in the Supplemental Materials & Methods.

## Data, Materials, and Software Availability

Code for simulations, data analysis, and data visualization have been deposited on GitHub (https://github.com/albobson/polio-res-eco-evo).

## Acknowledgments

We thank Feder and Kerr lab members and two reviewers for useful project and manuscript feedback. We thank Gaël Thebaud for the helpful advice on fitting the pocapavir fitness function. This work was possible by funding from NIH training grant T32-GM136534-02 supporting A. J. R and by the Environmental Biology Division from the National Science Foundation (grant number 2142718 to B.K.).

## Author contributions

A.J.R, B.K and A.F.F. contributed to study conceptualization and design. A.J.R. performed the analysis. A.J.R, B.K and A.F.F. interpreted the results, wrote the original draft of the manuscript and contributed to review and editing of the manuscript.

## References

[1] Elsa Hansen et al. “Antibiotics can be used to contain drug-resistant bacteria by maintaining sufficiently large sensitive populations”. en. In: PLOS Biology 18.5 (May 2020), e3000713. issn: 1545-7885. doi: 10.1371/journal.pbio.3000713.

[2] Caroline Colijn and Ted Cohen. “How competition governs whether moderate or aggressive treatment minimizes antibiotic resistance”. In: eLife 4 (Sept. 2015). Ed. by Michael Doebeli, e10559. issn: 2050-084X. doi: 10.7554/eLife.10559.

[3] Raul Andino and Esteban Domingo. “Viral quasispecies”. In: Virology. 60th Anniversary Issue 479–480 (May 2015), pp. 46–51. issn: 0042-6822. doi: 10.1016/j.virol.2015.03.022.

[4] Claus O Wilke and Isabel S Novella. “Phenotypic mixing and hiding may contribute to memory in viral quasispecies”. In: BMC Microbiology 3 (June 2003), p. 11. issn: 1471-2180. doi: 10.1186/1471-2180-3-11.

[5] Claude Loverdo and James O. Lloyd-Smith. “Inter-Generational Phenotypic Mixing in Viral Evolution”. In: Evolution; international journal of organic evolution 67.6 (June 2013), pp. 1815–1822. issn: 0014-3820. doi: 10.1111/evo.12048.

[6] Elizabeth J Tanner et al. “Dominant drug targets suppress the emergence of antiviral resistance”. In: eLife 3 (Nov. 2014). Ed. by Wenhui Li, e03830. issn: 2050-084X. doi: 10.7554/eLife.03830.

[7] Asher Leeks et al. “Open questions in the social lives of viruses”. en. In: Journal of Evolutionary Biology 36.11 (2023), pp. 1551–1567. issn: 1420-9101. doi: 10.1111/jeb.14203.

[8] Marc S. Collett et al. “Antiviral Activity of Pocapavir in a Randomized, Blinded, Placebo-Controlled Human Oral Poliovirus Vaccine Challenge Model”. In: The Journal of Infectious Diseases 215.3 (Feb. 2017), pp. 335–343. issn: 0022-1899. doi: 10.1093/infdis/jiw542.

[9] Elizabeth J. Tanner, Karla A. Kirkegaard, and Leor S. Weinberger. “Exploiting Genetic Interference for Antiviral Therapy”. en. In: PLOS Genetics 12.5 (May 2016), e1005986. issn: 1553-7404. doi: 10.1371/journal.pgen.1005986.

[10] Diana V Kouiavskaia et al. “Immunological and Pathogenic Properties of Poliovirus Variants Selected for Resistance to Antiviral Drug V-073”. en. In: Antiviral Therapy 16.7 (Oct. 2011), pp. 999–1004. issn: 1359-6535. doi: 10.3851/IMP1838.

[11] Hong-Mei Liu et al. “Characterization of Poliovirus Variants Selected for Resistance to the Antiviral Compound V-073”. In: Antimicrobial Agents and Chemotherapy 56.11 (Nov. 2012), pp. 5568–5574. doi: 10.1128/AAC.00539-12.

[12] Julie Copelyn et al. “Clearance of Immunodeficiency-associated Vaccine-derived Poliovirus Infection With Pocapavir”. en-US. In: The Pediatric Infectious Disease Journal 39.5 (May 2020), p. 435. issn: 0891-3668. doi: 10.1097/INF.0000000000002584.

[13] Sanet Torres-Torres et al. “First Use of Investigational Antiviral Drug Pocapavir (V-073) for Treating Neonatal Enteroviral Sepsis”. en-US. In: The Pediatric Infectious Disease Journal 34.1 (Jan. 2015), p. 52. issn: 0891-3668. doi: 10.1097/INF.0000000000000497.

[14] Ashleigh S. Griffin and Asher Leeks. “Exploiting social traits for clinical applications in bacteria and viruses”. en. In: npj Antimicrobials and Resistance 3.1 (Mar. 2025), pp. 1–9. issn: 2731-8745. doi: 10.1038/s44259-025-00091-6.

[15] Leor S. Weinberger, David V. Schaffer, and Adam P. Arkin. “Theoretical Design of a Gene Therapy To Prevent AIDS but Not Human Immunodeficiency Virus Type 1 Infection”. In: Journal of Virology 77.18 (Sept. 2003), pp. 10028–10036. issn: 0022-538X. doi: 10.1128/JVI.77.18.10028-10036.2003.

[16] Vincent T. Metzger, James O. Lloyd-Smith, and Leor S. Weinberger. “Autonomous Targeting of Infectious Superspreaders Using Engineered Transmissible Therapies”. In: PLoS Computational Biology 7.3 (Mar. 2011), e1002015. issn: 1553-734X. doi: 10.1371/journal.pcbi.1002015.

[17] Fathima N Nagoor Pitchai et al. “Engineered deletions of HIV replicate conditionally to reduce disease in nonhuman primates”. In: Science 385.6709 (2024), eadn5866.

[18] Sonali Chaturvedi et al. “Identification of a therapeutic interfering particle—A single-dose SARS-CoV-2 antiviral intervention with a high barrier to resistance”. English. In: Cell 184.25 (Dec. 2021), 6022–6036.e18. issn: 0092-8674, 1097-4172. doi: 10.1016/j.cell.2021.11.004.

[19] Sonali Chaturvedi et al. “A single-administration therapeutic interfering particle reduces SARS-CoV-2 viral shedding and pathogenesis in hamsters”. In: Proceedings of the National Academy of Sciences 119.39 (Sept. 2022), e2204624119. doi: 10.1073/pnas.2204624119.

[20] Veronica V. Rezelj et al. “Defective viral genomes as therapeutic interfering particles against flavivirus infection in mammalian and mosquito hosts”. en. In: Nature Communications 12.1 (Apr. 2021), p. 2290. issn: 2041-1723. doi: 10.1038/s41467-021-22341-7.

[21] Lei Zhang et al. “Adaptive therapy: a tumor therapy strategy based on Darwinian evolution theory”. In: Critical Reviews in Oncology/Hematology 192 (Dec. 2023), p. 104192. issn: 1040-8428. doi: 10.1016/j.critrevonc.2023.104192.

[22] Robert A. Gatenby and Joel S. Brown. “The Evolution and Ecology of Resistance in Cancer Therapy”. In: Cold Spring Harbor Perspectives in Medicine 10.11 (Nov. 2020), a040972. issn: 2157-1422. doi: 10.1101/cshperspect.a040972.

[23] Olha Puhach, Benjamin Meyer, and Isabella Eckerle. “SARS-CoV-2 viral load and shedding kinetics”. en. In: Nature Reviews Microbiology 21.3 (Mar. 2023), pp. 147–161. issn: 1740-1534. doi: 10.1038/s41579-022-00822-w.

[24] Hitoshi Kawasuji et al. “Transmissibility of COVID-19 depends on the viral load around onset in adult and symptomatic patients”. en. In: PLOS ONE 15.12 (Dec. 2020), e0243597. issn: 1932-6203. doi: 10.1371/journal.pone.0243597.

[25] Robert W. Eisinger, Carl W. Dieffenbach, and Anthony S. Fauci. “HIV Viral Load and Transmissibility of HIV Infection: Undetectable Equals Untransmittable”. In: JAMA 321.5 (Feb. 2019), pp. 451–452. issn: 0098-7484. doi: 10.1001/jama.2018.21167.

[26] Anne Sicard et al. “The Strange Lifestyle of Multipartite Viruses”. en. In: PLOS Pathogens 12.11 (Nov. 2016), e1005819. issn: 1553-7374. doi: 10.1371/journal.ppat.1005819.

[27] Christopher B. Brooke et al. “Most Influenza A Virions Fail To Express at Least One Essential Viral Protein”. In: Journal of Virology 87.6 (Mar. 2013), pp. 3155–3162. doi: 10.1128/jvi.02284-12.

[28] Alistair B Russell, Cole Trapnell, and Jesse D Bloom. “Extreme heterogeneity of influenza virus infection in single cells”. In: Elife 7 (2018), e32303.

[29] DS Burke. “Recombination in HIV: an important viral evolutionary strategy.” In: Emerging Infectious Diseases 3.3 (1997), pp. 253–259. issn: 1080-6040. doi: 10.3201/eid0303.970301.

[30] Elena V Romero and Alison F Feder. “Elevated HIV Viral Load is Associated with Higher Recombination Rate In Vivo”. In: Molecular Biology and Evolution 41.1 (Jan. 2024), msad260. issn: 1537-1719. doi: 10.1093/molbev/msad260.

[31] Michael B Schulte et al. “Experimentally guided models reveal replication principles that shape the mutation distribution of RNA viruses”. In: eLife 4 (Jan. 2015). Ed. by Stephen P Goffs, e03753. issn: 2050-084X. doi: 10.7554/eLife.03753.

[32] Michael B. Schulte and Raul Andino. “Single-Cell Analysis Uncovers Extensive Biological Noise in Poliovirus Replication”. en. In: Journal of Virology 88.11 (June 2014). Ed. by S. Perlman, pp. 6205–6212. issn: 0022-538X, 1098-5514. doi: 10.1128/JVI.03539-13.

[33] Roberto Mateo, Claude M. Nagamine, and Karla Kirkegaard. “Suppression of Drug Resistance in Dengue Virus”. In: mBio 6.6 (Dec. 2015), 10.1128/mbio.01960–15. doi: 10.1128/mbio.01960-15.

[34] Michael Hunter and Diana Fusco. “Superinfection exclusion: A viral strategy with short-term benefits and long-term drawbacks”. en. In: PLOS Computational Biology 18.5 (May 2022), e1010125. issn: 1553-7358. doi: 10.1371/journal.pcbi.1010125.

[35] Adeline Kerviel, Mengyang Zhang, and Nihal Altan-Bonnet. “A New Infectious Unit: Extra-cellular Vesicles Carrying Virus Populations”. In: Annual Review of Cell and Developmental Biology 37.1 (2021), pp. 171–197. doi: 10.1146/annurev-cellbio-040621-032416.

[36] Ying-Han Chen et al. “Phosphatidylserine Vesicles Enable Efficient En Bloc Transmission of Enteroviruses”. In: Cell 160.4 (Feb. 2015), pp. 619–630. issn: 0092-8674. doi: 10.1016/j.cell.2015.01.032.

[37] Denise Egger and Kurt Bienz. “Intracellular location and translocation of silent and active poliovirus replication complexes”. In: Journal of General Virology 86.3 (2005), pp. 707–718. issn: 1465-2099. doi: 10.1099/vir.0.80442-0.

[38] Purnell W. Choppin and Kathryn V. Holmes. “Replication of SV5 RNA and the effects of superinfection with poliovirus”. In: Virology 33.3 (Nov. 1967), pp. 442–451. issn: 0042-6822. doi: 10.1016/0042-6822(67)90119-5.

[39] Elizabeth R. Aguilera et al. “Plaques Formed by Mutagenized Viral Populations Have Elevated Coinfection Frequencies”. In: mBio 8.2 (Mar. 2017), 10.1128/mbio.02020–16. doi: 10.1128/mbio.02020-16.

[40] Andrea K. Erickson et al. “Bacteria Facilitate Enteric Virus Co-infection of Mammalian Cells and Promote Genetic Recombination”. English. In: Cell Host & Microbe 23.1 (Jan. 2018), 77–88.e5. issn: 1931-3128. doi: 10.1016/j.chom.2017.11.007.

[41] Marion S. Freistadt, Gerardo Kaplan, and Vincent R. Racaniello. “Heterogeneous Expression of Poliovirus Receptor-Related Proteins in Human Cells and Tissues”. In: Molecular and Cellular Biology 10.11 (Nov. 1990), pp. 5700–5706. issn: null. doi: 10.1128/mcb.10.11.5700-5706.1990.

[42] Rosa Molfetta et al. “CD155: A Multi-Functional Molecule in Tumor Progression”. en. In: International Journal of Molecular Sciences 21.33 (Jan. 2020), p. 922. issn: 1422-0067. doi: 10.3390/ijms21030922.

[43] Alex Farrell et al. “Semi-infectious particles contribute substantially to influenza virus withinhost dynamics when infection is dominated by spatial structure”. In: Virus Evolution 9.1 (Mar. 2023), vead020. issn: 2057-1577. doi: 10.1093/ve/vead020.

[44] P. J. Buontempo et al. “SCH 48973: a potent, broad-spectrum, antienterovirus compound”. eng. In: Antimicrobial Agents and Chemotherapy 41.6 (June 1997), pp. 1220–1225. issn: 0066-4804. doi: 10.1128/AAC.41.6.1220.

[45] A Schlegel et al. “Cellular origin and ultrastructure of membranes induced during poliovirus infection”. In: Journal of Virology 70.10 (Oct. 1996), pp. 6576–6588. doi: 10.1128/jvi.70.10.6576-6588.1996.

[46] Reyes R. Novoa et al. “Virus factories: associations of cell organelles for viral replication and morphogenesis”. en. In: Biology of the Cell 97.2 (2005), pp. 147–172. issn: 1768-322X. doi: 10.1042/BC20040058.

[47] Selma Dahmane et al. “Membrane-assisted assembly and selective secretory autophagy of enteroviruses”. en. In: Nature Communications 13.1 (Oct. 2022), p. 5986. issn: 2041-1723. doi: 10.1038/s41467-022-33483-7.

[48] Claus O. Wilke. “Quasispecies theory in the context of population genetics”. In: BMC Evolutionary Biology 5.1 (Aug. 2005), p. 44. issn: 1471-2148. doi: 10.1186/1471-2148-5-44.

[49] Mario Santer and Hildegard Uecker. “Evolutionary rescue and drug resistance on multicopy plasmids”. In: Genetics 215.3 (2020), pp. 847–868.

[50] Lei Sun et al. “Effective polyploidy causes phenotypic delay and influences bacterial evolvability”. In: PLoS Biology 16.2 (2018), e2004644.

[51] John P. DeLong et al. “Towards an integrative view of virus phenotypes”. en. In: Nature Reviews Microbiology 20.2 (Feb. 2022), pp. 83–94. issn: 1740-1534. doi: 10.1038/s41579-021-00612-w.

[52] Iris Yousaf et al. “Brain tropism acquisition: The spatial dynamics and evolution of a measles virus collective infectious unit that drove lethal subacute sclerosing panencephalitis”. en. In: PLOS Pathogens 19.12 (Dec. 2023), e1011817. issn: 1553-7374. doi: 10.1371/journal.ppat.1011817.

[53] Katherine S Xue et al. “Cooperation between distinct viral variants promotes growth of H3N2 influenza in cell culture”. In: Elife 5 (2016), e13974.

[54] Yuta Shirogane et al. “Collective fusion activity determines neurotropism of an en bloc transmitted enveloped virus”. In: Science Advances 9.4 (2023), eadf3731.

[55] Celia Perales et al. “Insights into RNA Virus Mutant Spectrum and Lethal Mutagenesis Events: Replicative Interference and Complementation by Multiple Point Mutants”. In: Journal of Molecular Biology 369.4 (June 2007), pp. 985–1000. issn: 0022-2836. doi: 10.1016/j.jmb.2007.03.074.

[56] R Core Team. R: A Language and Environment for Statistical Computing. R Foundation for Statistical Computing. Vienna, Austria, 2024. url: https://www.R-project.org/.

[57] Hadley Wickham. ggplot2: Elegant Graphics for Data Analysis. Springer-Verlag New York, 2016. isbn: 978-3-319-24277-4. url: https://ggplot2.tidyverse.org.

