## Supplementary Information for "Intracellular interactions shape antiviral resistance outcomes in poliovirus via eco-evolutionary feedback"

### Supplemental Material

#### Supplemental Figures

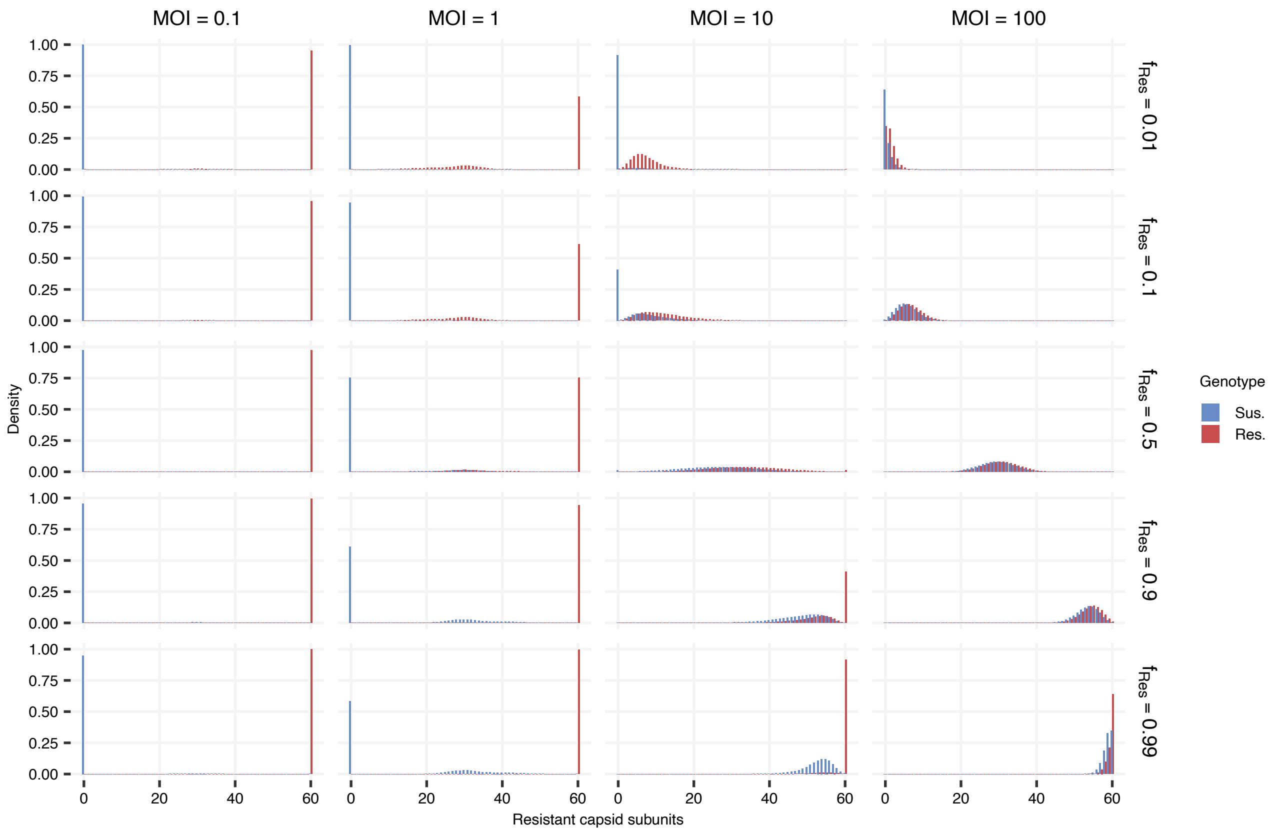


Figure S 1: **Distribution of capsid subunit compositions following viral replication.** For cellular populations infected by viruses with different frequencies of genotypic resistance ($f_{Res} \in \{0.01, 0.1, 0.5, 0.9, 0.99\}$, rows) and at different multiplicities of infection ($MOI \in \{0.1, 1, 10, 100\}$, columns), we plot the densities of virions emerging from these cells whose capsids contain different numbers of resistant capsid subunits. Outcomes are plotted separately based on if virions contain a susceptible genome (blue) or a resistant one (red). Simulations were run using standard model parameters reported in Table S1. At low MOIs (0.1 and 1), progeny genomes are predominantly encapsidated in capsids reflecting their own genotype (i.e., homogeneous capsids). At higher MOIs (10 and 100), progeny genomes are more frequently encapsidated in mixed capsids due to increased coinfection. Capsid composition is most mixed when resistant and susceptible genotypes are present at similar frequencies.

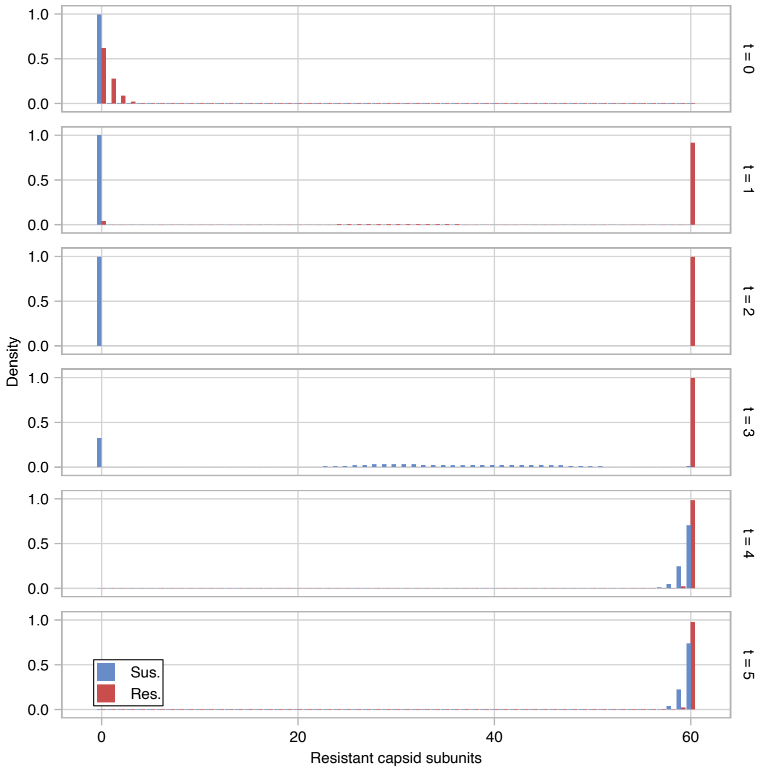


Figure S 2: **Distribution of capsid subunit compositions over time in a serial passaging experiment.** For each timestep in the serial passaging experiment shown in **Figure 4D** (rows), we plot the density of capsid subunit compositions (i.e., the number of resistant capsid subunits) for pre-neutralization virions based on whether they contain a susceptible (blue) or resistant (red) genome. The population was initialized with $f_{Res} = 10^{-4}$ and an MOI of 100. At $t = 0$, both resistant and susceptible genomes are packaged in highly susceptible capsids, but as the MOI drops, resistant and susceptible genomes are increasingly packaged in capsids matching their phenotypes. While MOI remains low and susceptible genomes become increasingly rare ($t = 3$), the phenotypic variance of susceptible genomes becomes large based on whether or not they coinfect with resistant ones. After rebound ($t \geq 4$), both resistant and susceptible genomes are packaged in highly resistant capsids.

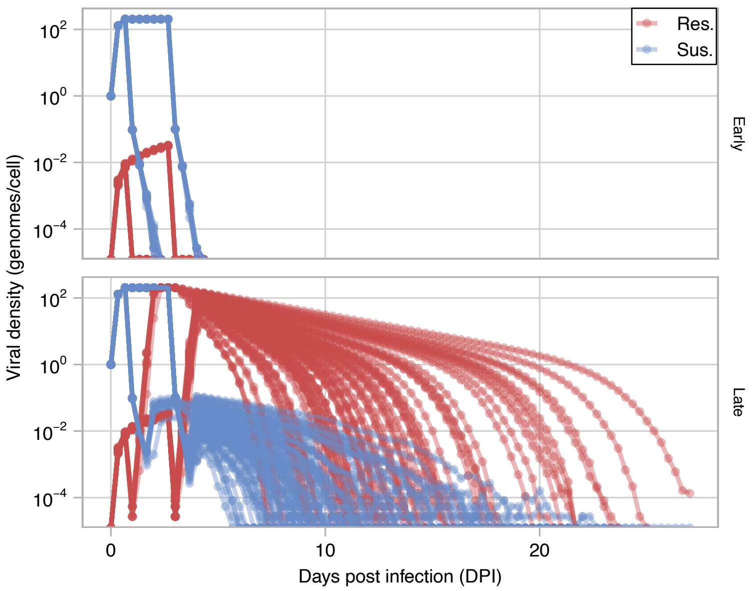


Figure S 3: **Population dynamics of individual simulations in the simulated pocapavir clinical trial.** For each simulated viral population in the pocapavir clinical trial summarized in **Figure 4F**, we plot the viral density (genomes/cell) for resistant (red) and susceptible (blue) genomes over time. We stratify populations into early clearers (top) or late clearers (bottom) based on if the clearance time was earlier that 7 days post infection (DPI) or later than or equal to 7 DPI. As in the clinical trial, simulations were administered pocapavir beginning either 24 or 72 hours post infection, which causes the two asynchronous drops in viral density 1 and 3 days post infection.

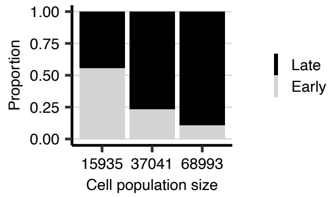


Figure S 4: **Smaller host cell population sizes lead to more frequent early clearance in simulated clinical trials.** Immune clearance parameters were optimized starting from three different initial host cell population sizes (15,000, 30,000, and 60,000) and converged to values similar to their starting conditions (15,935, 37,041, and 68,992, respectively). The frequency of early ($< 7$ days, black) versus late ($\geq 7$ days, grey) clearance was dependent on host cell population size, where early clearance was more common in simulations with smaller host cell population sizes ($n = 93$ simulations). In the main text, results are shown with the intermediate initial conditions ($\gamma = 37,041$).

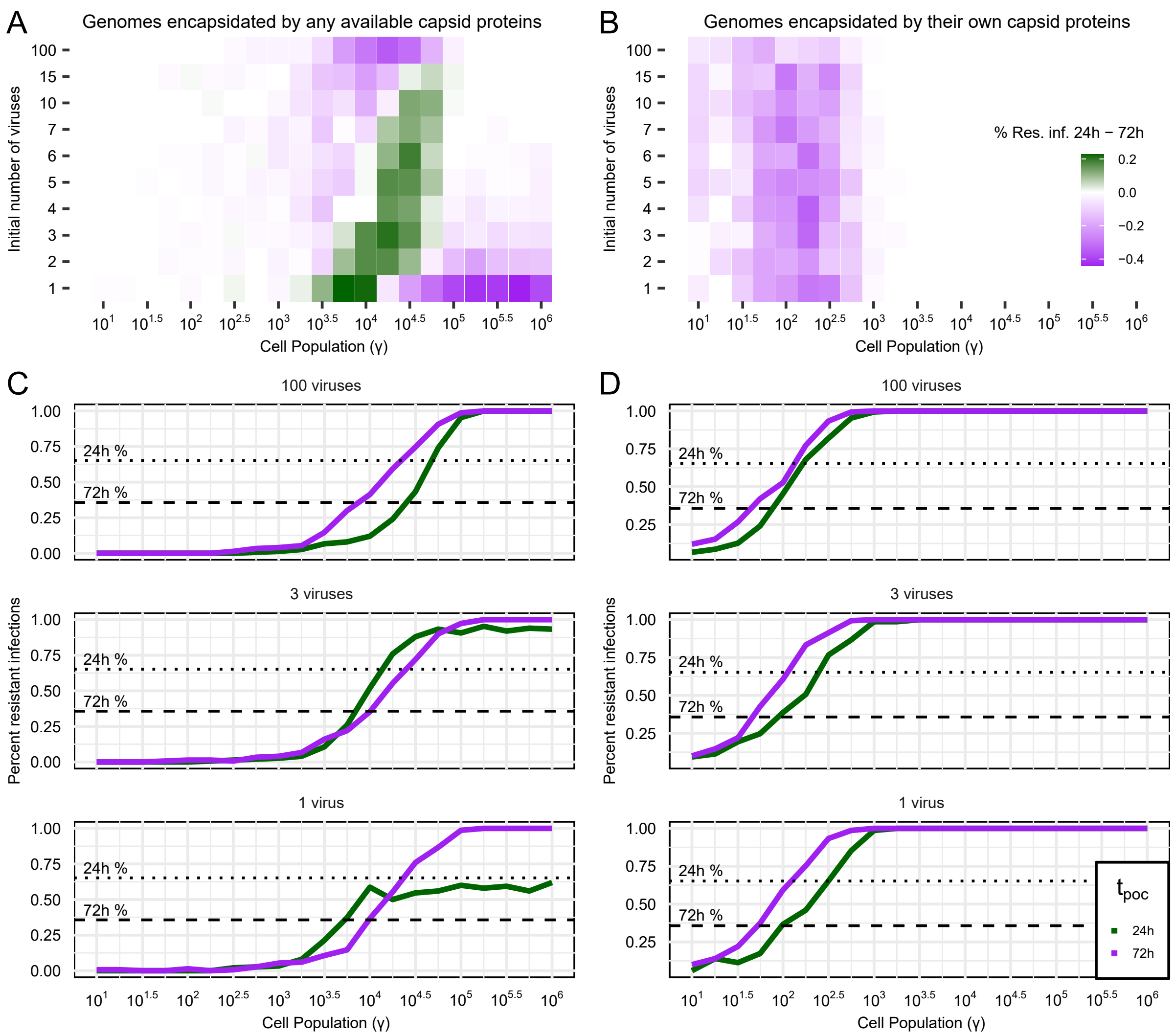


Figure S 5: **Intracellular protein sharing can increase the frequency of resistance evolution during early treatment.** Collett et al. [1] found that a greater proportion of viral populations treated 24 hours post-infection developed resistance than those treated 72 hours post-infection (15/23 versus 25/70; $p = 0.0163$, Fisher's exact test). We examined the emergence of this behavior using the model in this manuscript (in which genomes can be encapsidated by any available intracellular capsid proteins), and a model without intra-cellular mixing (in which genomes are encapsidated only by capsid proteins corresponding to their genotype). **(A, B)** We compared the rates of resistance evolution in the 24 and 72 hour treatment groups across host cell population sizes ($\gamma$) and initial numbers of infecting susceptible viruses ($n = 150$ per group and parameter set, $f_{Res} \geq 50\%$ defined as resistant). We identified conditions in which earlier treatment was associated with more resistance evolution in the 24 hour treatment group than the 72 hour treatment group (shown in green) in the intra-cellular mixing model **(A)** but not the non-mixing model **(B)**. **(C-D)** Example rows from **(A)** and **(B)** show the proportion of resistant infections in the 24 hour (green) versus 72 hour (purple) treatment groups under example initial conditions. Clinical resistance frequencies from Collett et al. [1] at the 24 hour and 72 hour treatment times are shown as dotted and dashed lines, respectively.

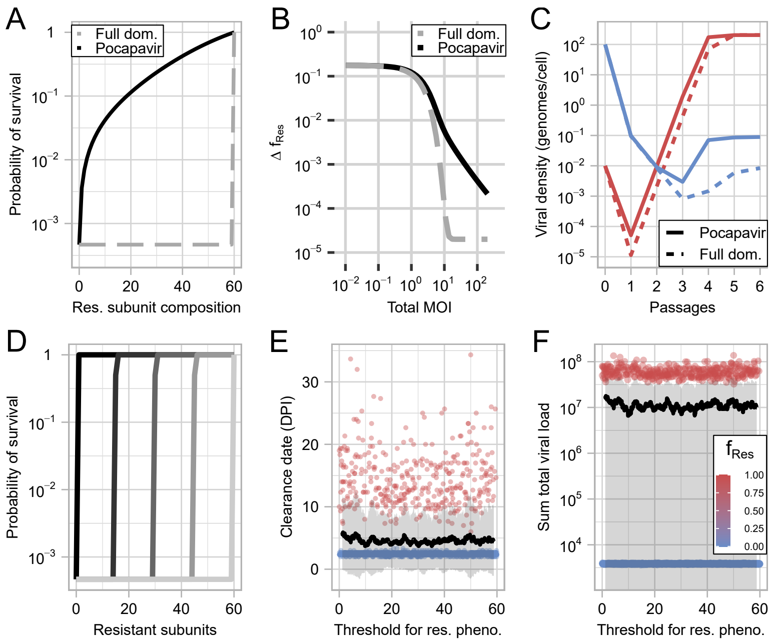


Figure S 6: **More "dominant" drugs do not suppress resistance better than pocapavir.** (**A**) We considered a hypothetical drug to which drug resistant capsid subunits only conferred a survival advantage if all 60 subunits were resistant (i.e., susceptible subunits fully dominate resistant ones). We compared this fully dominant drug (dashed grey) to pocapavir (solid black). (**B**) In single passage simulations assessing the change in resistance frequency ($\Delta f_{Res}$) as a function of the initial MOI, the fully dominant drug suppressed resistance marginally better than pocapavir at high MOI (initial $f_{Res} = 10^{-4}$). (**C**) However, in a deterministic serial passage experiment, both drugs (pocapavir, solid; the fully dominant drug, dashed) led to similar population trajectories for susceptible and resistant viral densities over time. In (**A-C**), we considered that a capsid must have 60 resistant subunits to have any increased survival probability. We next consider how lowering that threshold to generate a resistance response (i.e., 100% survival probability when treated with pocapavir) from 60 affects clinical outcomes. Five such thresholds are plotted in (**D**), but 2000 trials were run across a range of threshold values. Threshold values do not affect clearance date (**E**) or sum total viral load over the course of infection (**F**). Each point represents an individual simulation with a given resistant capsid threshold value to confer resistance, and points are colored by the frequency of genotypic resistance observed in the simulation over the course of infection. For both (**E**) and (**F**), the black line shows a rolling mean and the gray ribbon shows the variance around the mean.

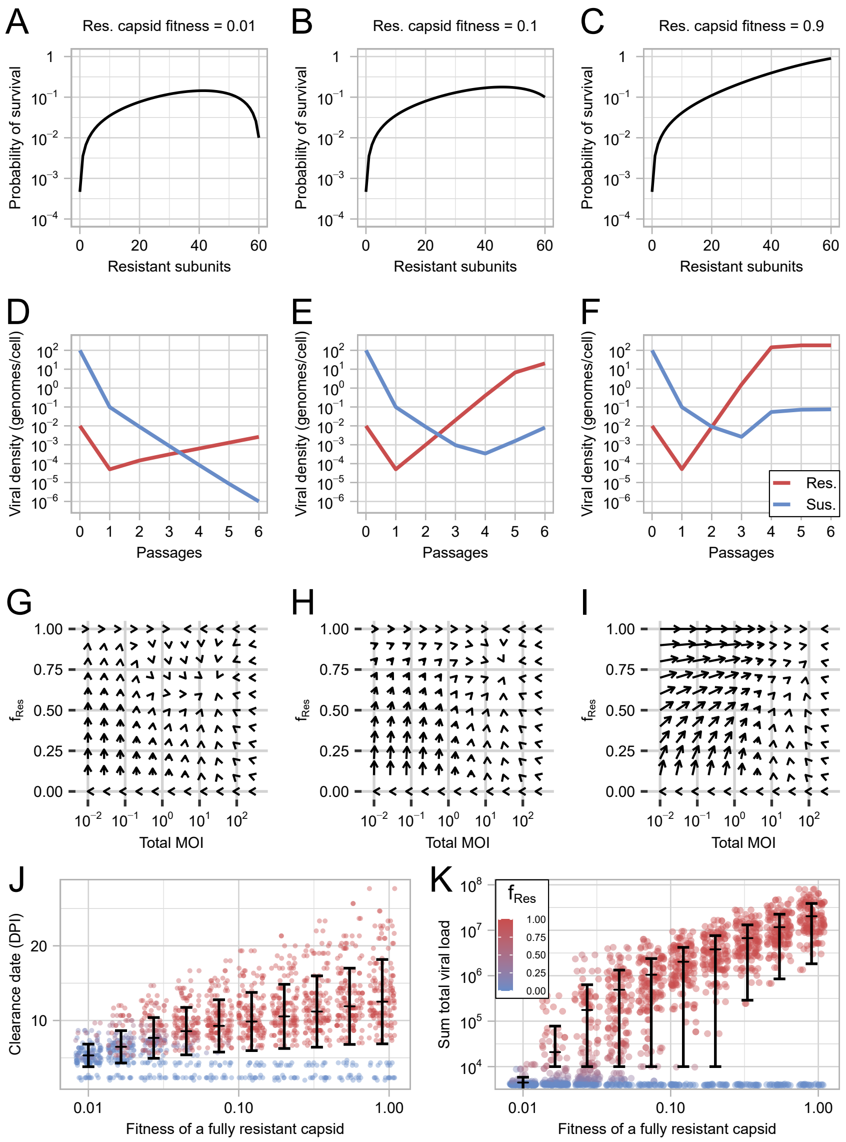


Figure S 7: **Fitness cost of phenotypic resistance impairs the spread of resistance and results in more favorable clinical outcomes.** (**A-C**) We generated compound fitness curves that accounted for both the standard fitness cost imposed by pocapavir and a linear fitness cost associated with greater numbers of resistant subunits per virion. The fitness of a fully resistant capsid is noted above its respective curve. Deterministic model simulations using the fitness functions in **A-C** reveal that greater costs of resistance slow the rate of resistance evolution and reduce viral densities over 6 passages (initial MOI = 100, initial $f_{Res} = 10^{-4}$, **D-F**) and reduce the equilibrium values of both total MOI and $f_{Res}$ (**G-I**). Clinical trials simulated with the same immune parameters and pocapavir treatment times as in Figure 4F, but incorporating fitness costs ($\kappa \in [0, 0.0165]$), showed earlier average clearance (**J**), reduced total viral loads (**K**), and lower resistance frequencies ($f_{Res}$; legend in **K** applies to point colors in **J**; horizontal line indicates mean value, bars show $\pm$ standard deviation).

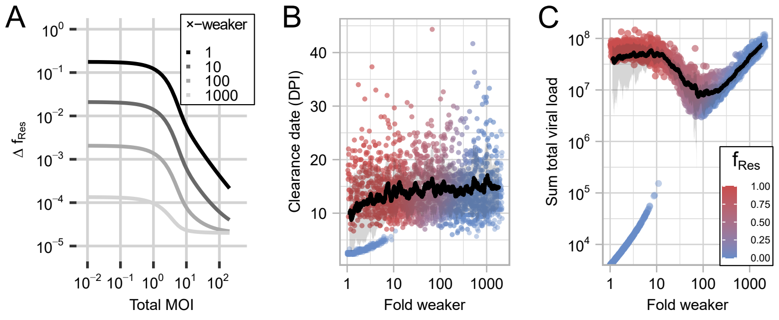


Figure S 8: **Reducing drug potency lowers selection for resistance and can maintain lower viral loads over time.** (**A**) In single passage simulations assessing the change in resistance frequency ($\Delta f_{Res}$) as a function of the initial MOI, less potent drugs (10×, 100×, 1000× weaker) led to smaller increases in the frequency of resistance than pocapavir (initial $f_{Res} = 10^{-4}$). Note, clinical trial outcomes for these specific drugs are plotted in **Figure 5**. We also ran 2000 clinical simulations exploring drugs with a range of potency reductions between 1× and 2147× weaker. (**B**) Drugs weaker than pocapavir had later clearance times due to the reduced probability of population bottlenecks leading to early extinction. (**C**) However, we observed a non-monotonic relationship between drug weakness and the sum total viral load, in which drugs with intermediate weakness (≈ 100× weaker) best reduce sum total viral load. For both (**B**) and (**C**), each point represents an individual simulation with a given resistant capsid threshold value to confer resistance, and points are colored by the frequency of genotypic resistance observed in the simulation over the course of infection (legend in (**C**)). The black line shows a rolling mean and the gray ribbon shows the variance around the mean.

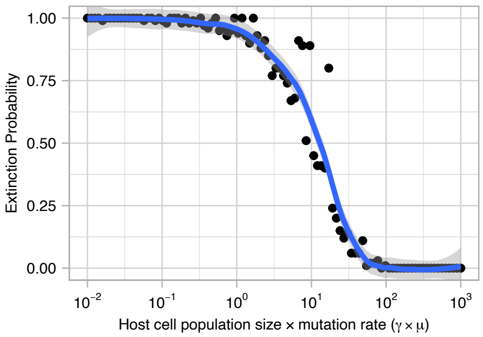


Figure S 9: **Stochastic extinction probability is determined by the product of the host cell population size and the mutation rate ($\gamma \times \mu$).** The relationship between stochastic extinction and $\gamma \times \mu$ was investigated using the stochastic version of the model. A total of 10,000 simulations were run across 100 different $\gamma \times \mu$ values (ranging from $10^{-2}$ to $10^{3}$, MOI = 1, $f_{\mathrm{Res}} = 0$). We plot the proportion of simulations in which the viral population went extinct within nine replication cycles. A larger host cell population ($\gamma$) increases the opportunities for *de novo* resistant mutants to arise, thereby decreasing the likelihood of stochastic extinction in the presence of pocapavir. The inflection point occurs near $\gamma \times \mu \approx 13$. This empirical value represents the threshold where, on average, 50% of simulations will generate a sufficient number of resistant genomes to establish at least one persistent lineage, overcoming both initial stochastic loss and future phenotypic mixing with susceptible genomes. This simulated threshold is similar to the analytical approximation of $\gamma \times \mu \approx 8.53$ (see Supplemental Materials & Methods – *In vivo* cellular population size).

| Parameter (units) | Variable | Estimated Value | Empirical Comparison |
|---|---|---|---|
| Fitness function, steepness | $k$ | 0.061675 | - |
| Fitness function, inflection | $i_0$ | 60 | - |
| Effective burst size (infections particles/cell) | $\beta$ | 203.17 | 23.89 - 277.78 (particles/cell[2, 3] $\times$ PFU/particle [4]) |
| Clinical model, host cell pop. (cells) | $\gamma$ | $37,041$ | - |
| Imm. clearance delay (hours) | $t_{\text{imm}}$ | 80 | 72 to 125 [5] |
| Imm. clearance distribution, mean | $\xi$ | $-1.424$ | - |
| Imm. clearance distribution, SD | $\tau$ | 0.460 | - |

##### Parameter Estimates

Table S 1: Estimated model parameters and their associated notation, estimated values and empirical comparisons. Parameter estimates were obtained by fitting the model to cell culture and clinical trial data. In brief, burst size and fitness function parameters were estimated by fitting to cell culture data reported in Tanner et al. [6], and host cell population and immune clearance parameters were estimated by fitting to clinical data from a placebo-treated group reported by Collett et al. [1] (see Materials & Methods—Parameter inference).

### Supplemental Materials and Methods

#### Overview

Here, we describe each step of the model using relevant probability distributions. Random variables, dynamic processes that involve stochasticity, are denoted by uppercase Roman letters with subscripts. Specific dynamic values are represented by lowercase Roman letters with corresponding subscripts. Input model parameters, whether inferred or drawn from the literature, are denoted by Greek letters and held constant across individual runs of a model. For conciseness, we describe the methods in terms of the stochastic model. The deterministic version follows the same probability distributions but accounts for each possible outcome (i.e., integrates over the probability distribution) instead of drawing specific values.

All simulations and analyses were performed in `R`, edition 4.1.0 [7]. Data visualization was performed with the `ggplot2` package [8].

#### Poliovirus replication model

Poliovirus replication is simulated as a discrete-time dynamical model that tracks abundances of viable resistant and susceptible poliovirus genomes after each round of viral replication, the distribution of virus capsid subunit compositions, and the number of infected and uninfected host cells. A single round of replication proceeds in the following steps:

##### Viral entry

We assume that cells can be infected by multiple viruses (i.e., that there is no superinfection exclusion), and that the probability that a cell is infected by a given virus is uniform across the cell population (i.e., there is no preferential infection). We let $r_{\text{tot},t}$ and $s_{\text{tot},t}$ be the number of virions respectively carrying resistant and susceptible viral genomes in the viral population at a given time, $t$. The total number of genomes are:

$$v_{\text{tot},t} = r_{\text{tot},t} + s_{\text{tot},t}.$$

We let $\gamma$ be the total number of host cells. The multiplicity of infection (MOI) is defined by the following ratio $m$:

$$m = \frac{v_{\text{tot},t}}{\gamma}.$$

We note that $m$ is undefined if $\gamma = 0$. Unless otherwise specified, the host cell population was held at $2.9 \times 10^6$ cells per mL, in accordance with the cell density in HeLa cultures. The MOI is 0 when $v_{\text{tot}} = 0$, and the simulation is terminated when that occurs.

We assume that virions do not interfere with other virions in terms of binding and entry into host cells. The number of genotypically resistant viruses infecting a given cell is a random variable, $R_{\text{inf}}$, where:

$$\Pr(R_{\text{inf}} = r_{\text{inf}}) = \binom{r_{\text{tot},t}}{r_{\text{inf}}} \left(\frac{1}{\gamma}\right)^{r_{\text{inf}}} \left(1 - \frac{1}{\gamma}\right)^{r_{\text{tot},t} - r_{\text{inf}}}.$$

That is, given $r_{\text{tot},t}$, $R_{\text{inf}} \sim \text{Bin}\left(r_{\text{tot},t}, \frac{1}{\gamma}\right)$. We note that if both $\gamma$ and $r_{\text{tot},t}$ are large, $\text{Bin}\left(r_{\text{tot},t}, \frac{1}{\gamma}\right) \approx \text{Pois}(\frac{r_{\text{tot},t}}{\gamma})$, which is how viral entry is often modeled.

The same relationship holds for the random variable $S_{\text{inf}}$, a random variable representing the number of susceptible viruses that infect a given cell (i.e., $S_{\text{inf}} \sim \text{Bin}\left(s_{\text{tot},t}, \frac{1}{\gamma}\right)$). The total number of viruses that have infected a given cell, $v_{\text{inf}}$, can be described by the equation: $v_{\text{inf}} = r_{\text{inf}} + s_{\text{inf}}$.

In the deterministic version of the model, $r_{\text{tot},t}$ and $s_{\text{tot},t}$ may take on non-integer values, making a binomial distribution ill-defined (as the number of trials is not an integer). We illustrate our solution to this problem by outlining how we derive the probability that a cell is infected by a certain number of resistant genomes, given a non-integer value of $r_{\text{tot},t}$.

Let $r_{\text{tot},t} = z + f$, where $z$ is a non-negative integer and $f$ is a real-valued number strictly between 0 and 1. For a given cell, we define the probability that there are $r_{\text{inf}}$ infecting viruses (with resistant genomes) to be:

$$\Pr(r_{\text{inf}}) = \begin{cases} (1-f)\binom{z}{r_{\text{inf}}} \left(\frac{1}{\gamma}\right)^{r_{\text{inf}}} \left(1 - \frac{1}{\gamma}\right)^{z - r_{\text{inf}}} + f\binom{z+1}{r_{\text{inf}}} \left(\frac{1}{\gamma}\right)^{r_{\text{inf}}} \left(1 - \frac{1}{\gamma}\right)^{z+1-r_{\text{inf}}} & \text{if } 0 \leq r_{\text{inf}} \leq z \\ f\left(\frac{1}{\gamma}\right)^{z+1} & \text{if } r_{\text{inf}} = z+1. \end{cases}$$

Letting $R_{\text{inf}}$ be the relevant random variable, we note that:

$$\mathbb{E}[R_{\text{inf}}] = \frac{z+f}{\gamma} = \frac{r_{\text{tot},t}}{\gamma},$$

which is MOI for viruses with resistant genomes at time $t$.

##### Genome replication

The number of new replicated genomes that an individual cell produces is a random variable $V_{\text{rep}}$. We let $\beta$ be the average number of infectious viral particles released per cell (i.e., the effective burst size). The number of replicated genomes per cell is a random variable, $V_{\text{rep}}$, where:

$$\Pr(V_{\text{rep}} = v_{\text{rep}} \mid v_{\text{inf}}) = \begin{cases} \dfrac{\beta^{v_{\text{rep}}} e^{-\beta}}{v_{\text{rep}}!}, & \text{if } v_{\text{inf}} > 0, \\ 0, & \text{if } v_{\text{inf}} = 0. \end{cases}$$

That is, given $v_{\text{inf}} > 0$, $V_{\text{rep}} \sim \text{Pois}(\beta)$.

We assume that resistant and susceptible genomes replicate at the same rate. The number of replicated resistant genomes inside of the cell, before considering mutation, is the random variable $R_{\text{rep}}$. The probability that a resistant genome replicates is defined by the ratio $r_{\text{inf}}/v_{\text{inf}}$, the frequency of resistant genomes upon initial infection. The probability that $R_{\text{rep}}$ takes on a specific value is:

$$\Pr\Big(R_{\text{rep}} = r_{\text{rep}} \mid \{v_{\text{rep}}, v_{\text{inf}}, r_{\text{inf}}\}\Big) = \binom{v_{\text{rep}}}{r_{\text{rep}}} \left(\frac{r_{\text{inf}}}{v_{\text{inf}}}\right)^{r_{\text{rep}}} \left(\frac{v_{\text{inf}} - r_{\text{inf}}}{v_{\text{inf}}}\right)^{v_{\text{rep}} - r_{\text{rep}}}.$$

That is, given $v_{\text{rep}}, v_{\text{inf}}$, and $r_{\text{inf}}$, $R_{\text{rep}} \sim \text{Bin}\Big(v_{\text{rep}}, \frac{r_{\text{inf}}}{v_{\text{inf}}}\Big)$.

In our model, the number of replicated susceptible genomes inside of a cell, $s_{\text{rep}}$, is defined by the equation:

$$s_{\text{rep}} = v_{\text{rep}} - r_{\text{rep}}.$$

##### Genome mutation

For simplicity, we assume that the same infecting templates are used for each progeny genome, and thus newly replicated genomes are not used in further replication. We assume that susceptible genomes can mutate during replication to become resistant and likewise, resistant genomes can mutate to become susceptible. Per replication event, the probability of mutation from a susceptible to a resistant genotype or from a resistant to a susceptible genotype is $\mu$. The number of new mutant resistant and susceptible viral genomes in a cell are the random variables $R_{\text{mut}}$ and $S_{\text{mut}}$. Given the number of resistant and susceptible genomes in the cell, the probabilities that these random variables take on specific values can be described by:

$$\Pr\Big(R_{\text{mut}} = r_{\text{mut}} \mid s_{\text{rep}}\Big) = \binom{s_{\text{rep}}}{r_{\text{mut}}} \mu^{r_{\text{mut}}} (1-\mu)^{s_{\text{rep}}-r_{\text{mut}}},$$

$$\Pr\Big(S_{\text{mut}} = s_{\text{mut}} \mid r_{\text{rep}}\Big) = \binom{r_{\text{rep}}}{s_{\text{mut}}} \mu^{s_{\text{mut}}} (1-\mu)^{r_{\text{rep}}-s_{\text{mut}}}.$$

That is, given $s_{\text{rep}}$, $R_{\text{mut}} \sim \text{Bin}(s_{\text{rep}}, \mu)$ and given $r_{\text{rep}}$, $S_{\text{mut}} \sim \text{Bin}(r_{\text{rep}}, \mu)$.

We calculated $\mu$ by determining the probability that a newly synthesized poliovirus genome would contain one of the two most common single amino acid substitutions that confer resistance to pocapavir (I194M or A24V) [9, 10]. We note that other amino acid substitutions that confer resistance to pocapavir have been identified, however, they are either found at low frequency in cell culture, and/or require transversion mutations which are 10× to 100× rarer than transition mutations [11], and are therefore not considered here. Both I194M and A24V can be generated by single transition mutations, so we used the per-replication event mutation rate of $\mu = 2 \times 10^{-5}$ as described by Schulte et al. [12] for our mutation rate.

The total number of resistant and susceptible genomes inside of the cell after replication and mutation are $r_{\text{pool}}$ and $s_{\text{pool}}$. Given $r_{\text{rep}}$, $s_{\text{rep}}$, $r_{\text{mut}}$ and $s_{\text{mut}}$, they are defined by:

$$r_{\text{pool}} = r_{\text{rep}} - s_{\text{mut}} + r_{\text{mut}},$$

$$s_{\text{pool}} = s_{\text{rep}} - r_{\text{mut}} + s_{\text{mut}}.$$

##### Capsid formation

We assume that both resistant and susceptible genotypes produce their corresponding capsid subunits at equal rates, and that newly replicated genomes do not contribute to capsid subunit creation. Thus, the frequency of resistant subunits in the pool of available subunits is $r_{\text{inf}}/v_{\text{inf}}$. Each newly-replicated genome inside of the cell draws from the available pool of subunits to build its capsid. We let $\sigma$ represent the number of subunits of which a capsid is composed. We assume that the spatial distribution of each subunit within the capsid has no effect on the capsid's fitness, and therefore we describe viral phenotype as simply the number of resistant subunits ($i$) out of $\sigma$. The number of resistant capsid subunits in an individual virion is a random variable $I$. The probability that $I = i$ is $p_i$. This probability is:

$$p_i = \Pr(I = i \mid \{v_{\text{inf}}, r_{\text{inf}}\}) = \binom{\sigma}{i} \left(\frac{r_{\text{inf}}}{v_{\text{inf}}}\right)^i \left(1 - \frac{r_{\text{inf}}}{v_{\text{inf}}}\right)^{\sigma - i}.$$

That is, given $v_{\text{inf}}$ and $r_{\text{inf}}$, $I \sim \text{Bin}\left(\sigma, \frac{r_{\text{inf}}}{v_{\text{inf}}}\right)$. We note that when $v_{\text{inf}} = 0$, $I$ is undefined. There are a total of 60 pocapavir binding sites per poliovirus virion, thus, in our simulations, $\sigma = 60$.

The number of resistant and susceptible genomes, after replication and mutation, that are packaged into capsids with $I = i$ resistant subunits are the multivariate random variables $R_{\text{pack}}$ and $S_{\text{pack}}$, where the $i^{th}$ components are $R_{\text{pack},i}$ and $S_{\text{pack},i}$. The probabilities that $R_{\text{pack},i}$ and $S_{\text{pack},i}$ take on values $r_{\text{pack},i}$ and $s_{\text{pack},i}$ are, respectively:

$$\Pr\Big(R_{\text{pack},0} = r_{\text{pack},0}, \ldots, R_{\text{pack},\sigma} = r_{\text{pack},\sigma} \mid r_{\text{pool}}\Big) = \frac{r_{\text{pool}}!}{r_{\text{pack},0}!\, r_{\text{pack},1}! \cdots r_{\text{pack},\sigma}!}\, p_0^{r_{\text{pack},0}}\, p_1^{r_{\text{pack},1}} \cdots p_\sigma^{r_{\text{pack},\sigma}};$$

$$\Pr\Big(S_{\text{pack},0} = s_{\text{pack},0}, \ldots, S_{\text{pack},\sigma} = s_{\text{pack},\sigma} \mid s_{\text{pool}}\Big) = \frac{s_{\text{pool}}!}{s_{\text{pack},0}!\, s_{\text{pack},1}! \cdots s_{\text{pack},\sigma}!}\, p_0^{s_{\text{pack},0}}\, p_1^{s_{\text{pack},1}} \cdots p_\sigma^{s_{\text{pack},\sigma}}.$$

That is, given $r_{\text{pool}}$, $R_{\text{pack}} \sim \text{Multinom}(r_{\text{pool}}, p_0, \ldots, p_\sigma)$ and given $s_{\text{pool}}$, $S_{\text{pack}} \sim \text{Multinom}(s_{\text{pool}}, p_0, \ldots, p_\sigma)$. At this point, the viral output from all cells are organized into bins according to subunit phenotype and genotype.

##### Pocapavir neutralization model

We assume that pocapavir molecules exist in excess compared to the number of susceptible subunits in the population and that the spatial distribution of resistant capsid subunits within a capsid does not affect drug binding and subsequent fitness of the virion. We let $\omega_i$ be the probability that a capsid with $i$ resistant capsid subunits survives, specifically due to the presence or absence of pocapavir. We assume that all infectious virions survive in the absence of pocapavir (i.e., without drug, $\omega_i = 1$ for all $i$).

We used cell culture data from Tanner et al. [6] to estimate the probability of a virion surviving pocapavir, given that it has $i$ resistant capsid subunits. Their study measured survival rates of pure susceptible and pure resistant poliovirus cultures in the presence of pocapavir. They also examined mixed cultures, fixing the MOI of resistant viruses at 10 while varying the MOI of susceptible viruses from 0 to 100 [6].

We make two assumptions to estimate the probability of survival given a virion's capsid phenotype composition: 1) the average progeny virion's capsid contains resistant subunits in the same ratio as the population's initial ratio of resistant to susceptible genomes; 2) dividing an individual culture's survival by the maximum potential survival (the survival of the pure resistant strain) provides an accurate estimate for the survival of the average capsid phenotype.

With these assumptions, we generated a set of survival probabilities across the range of possible capsid phenotypes, from 0 to 60 resistant subunits (the maximum number of binding sites) (**Figure 3 B**). We have accurate estimates for both ends of the phenotypic distribution, as pure cultures should produce progeny with capsids that are entirely resistant or susceptible. However, a distribution of capsid phenotypes should exist in the mixed cell cultures, as progeny phenotype is dependent on the initial infecting genomes. For this reason, we fit a logistic function through the data, while holding the probabilities of survival from pure culture data fixed. Let $L(i)$ be the standard form of the logistic function:

$$L(i) = \frac{1}{1 + e^{-k(i-i_0)}}$$

where $i$ is the number of resistant subunits, $i_0$ is the inflection point of the curve, and $k$ is the steepness of the curve.

Let $\omega(i,t)$ be the survival probability of a capsid composed of $i$ resistant subunits at time $t$. Pocapavir is administered after a specified number of viral replications, $t_{\text{poc}}$. If $t < t_{\text{poc}}$, a capsid has a pocapavir survival probability of 1, regardless of capsid subunit composition. If $t \geq t_{\text{poc}}$, the survival probability is described by the equation:

$$\omega(i, t \geq t_{poc}) = y_0 + (y_\sigma - y_0) \cdot \frac{L(i) - L(0)}{L(\sigma) - L(0)}$$

where $y_0$ is the survival probability of a capsid with zero resistant subunits, and $y_\sigma$ is the survival probability of a capsid with $\sigma$ resistant subunits (a fully resistant capsid). This equation ensures that when $i = 0$, $\omega(0, t \geq t_{poc}) = y_0$, and likewise, when $i = 60$, $\omega(60, t \geq t_{poc}) = y_\sigma$. The values for $i_0$ and $k$ were then inferred by nonlinear least squares regression (see Materials & Methods—Parameter inference).

Taken together, the survival probability of a capsid with $i$ resistant subunits at time $t$ is described by the set of equations:

$$\omega(i,t) = \begin{cases} 1, & \text{if } t < t_{\text{poc}}, \\ \\ y_0 + (y_\sigma - y_0) \cdot \frac{L(i)-L(0)}{L(\sigma)-L(0)}, & \text{if } t \geq t_{\text{poc}}. \end{cases}$$

The number of capsids carrying resistant or susceptible genomes that survive pocapavir and are composed of $i$ resistant subunits are random numbers, $R_{\text{surv},i}$ and $S_{\text{surv},i}$. The probability that $R_{\text{surv},i}$ and $S_{\text{surv},i}$ take on specific values are described by:

$$\Pr\Big(R_{\text{surv},i} = r_{\text{surv},i} \mid r_{\text{pack},i}\Big) = \binom{r_{\text{pack},i}}{r_{\text{surv},i}} \, \omega(i,t)^{r_{\text{surv},i}} \, (1 - \omega(i,t))^{r_{\text{pack},i} - r_{\text{surv},i}},$$

$$\Pr\Big(S_{\text{surv},i} = s_{\text{surv},i} \mid s_{\text{pack},i}\Big) = \binom{s_{\text{pack},i}}{s_{\text{surv},i}} \, \omega(i,t)^{s_{\text{surv},i}} \, (1 - \omega(i,t))^{s_{\text{pack},i} - s_{\text{surv},i}}.$$

Therefore, given $r_{\text{pack},i}$, $R_{\text{surv},i} \sim \text{Bin}(r_{\text{pack},i}, \omega(i,t))$. Similarly, given $s_{\text{pack},i}$, $S_{\text{surv},i} \sim \text{Bin}(s_{\text{pack},i}, \omega(i,t))$.

Unless otherwise noted, we assume that subunit composition does not affect infectivity, nor future immune survival. For this reason, the total number of infectious virions carrying resistant or susceptible genomes after replication is complete is the sum of all $i$ subunit states for each respective random variable. We let $r_{\text{sum}}$ and $s_{\text{sum}}$ represent the sum of all resistant and susceptible genotypes that survive pocapavir. They are defined by:

$$r_{\text{sum}} = \sum_{i=0}^{\sigma} r_{\text{surv},i},$$

$$s_{\text{sum}} = \sum_{i=0}^{\sigma} s_{\text{surv},i}.$$

#### Immune neutralization

Similar to previous models of host immune clearance during viral infection [13], we model immune clearance as an exponentially decreasing probability of virion survival, with a tunable delay in immune activation. Let $t_{\text{imm}}$ represent the delay before immune clearance begins. Once $t > t_{\text{imm}}$, the probability of a virion surviving immune clearance at each time step depends on a random variable representing initial immune sensitivity, $I_{\text{init}}$, which is drawn at the beginning of each simulation (and therefore, $I_{\text{init}}$ is not time dependent). The probability that $I_{\text{init}}$ takes on a specific value is:

$$\Pr(I_{\text{init}} = i_{\text{init}}) = \frac{1}{i_{\text{init}}\sqrt{2\pi\tau^2}} e^{-\frac{(\ln(i_{\text{init}})-\xi)^2}{2\tau^2}}.$$

That is, $I_{\text{init}} \sim \text{Lognormal}(\xi, \tau^2)$, where $\xi$ and $\tau^2$ are the distribution's mean and variance, respectively, and are inferred from the pocapavir clinical trial placebo group clearance times (see Materials & Methods—Parameter inference).

Let $d(t)$ be a function that describes the probability that a virion survives the immune system at time $t$. This function is described by:

$$d(t) = \begin{cases} \exp\left(-i_{\text{init}}(t - t_{\text{imm}})\right) & \text{if } t \geq t_{\text{imm}}, \\ 1 & \text{if } t < t_{\text{imm}}. \end{cases}$$

The number of virions that survive immune clearance and carry resistant or susceptible genomes are random variables $R_{\text{imm}}$ and $S_{\text{imm}}$, respectively. The probabilities that $R_{\text{imm}}$ and $S_{\text{imm}}$ take on specific values are described by:

$$\Pr\Big(R_{\text{imm}} = r_{\text{imm}} \mid r_{\text{sum}}\Big) = \binom{r_{\text{sum}}}{r_{\text{imm}}} d(t)^{r_{\text{imm}}} \Big(1 - d(t)\Big)^{r_{\text{sum}} - r_{\text{imm}}},$$

$$\Pr\Big(S_{\text{imm}} = s_{\text{imm}} \mid s_{\text{sum}}\Big) = \binom{s_{\text{sum}}}{s_{\text{imm}}} d(t)^{s_{\text{imm}}} \Big(1 - d(t)\Big)^{s_{\text{sum}} - s_{\text{imm}}}.$$

And so, given $r_{\text{sum}}$ and $s_{\text{sum}}$, $R_{\text{imm}} \sim \text{Bin}(r_{\text{sum}}, d(t))$ and $S_{\text{imm}} \sim \text{Bin}(s_{\text{sum}}, d(t))$. This is the last step in the replication cycle. The total resistant and susceptible viral population sizes after the replication cycle (i.e., the values for the next timestep) are $r_{\text{tot},t+1} = r_{\text{imm}}$ and $s_{\text{tot},t+1} = s_{\text{imm}}$. Note, immune clearance is considered in the stochastic model only.

#### Variation in dominance of the susceptible phenotype

We define the dominance of a discrete, multi-component phenotype as the relationship between structural composition (here, in the form of the number of susceptible/resistant subunits that make up a capsid) and phenotypic outcome (here, survival in the presence of pocapavir). For example, if a single resistant subunit conferred complete survival, resistance would be fully dominant. Alternatively, if capsids needed to be composed entirely of resistant subunits for a virion to escape neutralization, and were otherwise neutralized at the same rate as a fully susceptible capsid, resistance would be fully recessive.

To explore the impact of susceptible dominance over the resistant phenotype, we varied the pocapavir fitness function's parameters to simulate different dominance relationships. For simplicity, we assumed a step-wise threshold number of resistant subunits at or above which which a virion is fully resistant to drug neutralization. Using the logistic function as our base, we used a steepness coefficient of $k = 100$ to simulate a step-like function, and varied $i_0$ to to set the inflection point, corresponding to the minimum number of resistant subunits needed to render a virion resistant to drug.

To assess the full range of resistant subunit thresholds (i.e., the number of resistant subunits required to render a virion fully resistant to the drug), we used 2,000 hypothetical curves, with each $i_0$ split evenly between 0 and 60 (the maximum number of subunits in a poliovirus virion). We then simulated one clinical run (stochastic simulations incorporating immune clearance) for each of these neutralization curves. The only change that we made to these simulations was in the pocapavir neutralization step and all parameters were otherwise the same between runs. We calculated the rolling mean and standard deviation of clinical outcomes (clearance time and total viral load) using the functions `rollapply()` and `rollmean()` from the `zoo` package [14].

##### Fitness cost of resistance

We assumed that a fitness cost of resistance for a modified capsid subunit would occur extracellularly on capsid phenotypes rather than intracellularly at genome replication. We therefore encoded a fitness cost through a linear function in which each resistant capsid subunit reduced a virion's survival probability by a fixed amount, $\kappa$, such that a virion with with $i$ resistant subunits ($i \in [0, \sigma]$) had survival probability $K(i)$, where:

$$K(i) = 1 - \kappa \times i.$$

We imposed the fitness cost at the same extracellular survival step as pocapavir-mediated neutralization, by multiplying $\omega(i,t)K(i)$ element-wise before binomially sampling. While the expected fitness cost of pocapavir resistance mutations is $\kappa = 0$, we tested both deterministic and stochastic model outcomes for $\kappa \in [0.0165, 0]$. Throughout the manuscript, $\kappa = 0$ unless otherwise noted.

##### Variation in drug potency

We define drug potency as the ability of the drug to neutralize viruses for some capsid subunit composition. That is, a drug less potent than pocapavir would allow more viruses to survive treatment. We generated hypothetical, less potent drug survival probabilities by setting the minimum survival probability to a desired minimum survival value, $y_0'$, and re-scaling the pocapavir survival probabilities to match the overall relationship with the new minimum fitness value (**Figure 5** ). We define a new function, $\omega'(i,t)$, as a rescaled, less potent drug, which maintains the overall shape of the survival curve for capsids in the presence of pocapavir. This rescaled function is also time-dependent, and takes the form:

$$\omega'(i,t) = \begin{cases} 1, & \text{if } t < t_{\text{poc}}, \\ \\ y'_0 + \frac{(\omega(i,t)-y_0)(y_\sigma - y'_0)}{y_\sigma - y_0}, & \text{if } t \geq t_{\text{poc}}, \end{cases}$$

where $y'_0$ is the new fitness value of a capsid entirely composed of susceptible subunits, $y_0$ is the fitness value of a capsid entirely composed of susceptible subunits in the presence of pocapavir, and $y_\sigma$ is the fitness value of a capsid composed entirely of resistant subunits.

#### *In vivo* cellular population size

We estimated the *in vivo* cellular population size $\gamma$ jointly with the immune parameters as described in section Materials & Methods—Parameter inference. However, we found that initializing optimization with different starting values of $\gamma$ led to convergence at different fit $\gamma$ values that were similar to the starting values. This suggests that the clearance timings may not provide strong evidence that a particular *in vivo* population size is correct. We therefore report an intermediate value of $\gamma$ in the main text, but also present results with two other initial conditions in **Figure S4**. Choosing lower values of $\gamma$ increases the frequency of early viral extinction, while choosing higher values of $\gamma$ decreases its frequency.

The probability of a viral population avoiding extinction through drug resistance is fundamentally linked to the product of the total number of infected cells, $\gamma$, and the viral mutation rate, $\mu$. The establishment of a resistant population is a stochastic process that requires a newly generated resistant virion to first survive pocapavir neutralization, then successfully infect a host cell and propagate. Here, we focus on the initial, critical step where a *de novo* resistant genome, packaged within a susceptible capsid, must survive antiviral treatment.

The probability of a single phenotypically susceptible capsid surviving pocapavir is $y_0$. Consequently, the probability pocapavir-mediated neutralization is $1 - y_0$. For a fully susceptible population of viruses at an initial MOI = 1 infecting a population of $\gamma$ host cells, with each producing an effective viral burst of size $\beta$, the expected number of *de novo* resistant genomes packaged into phenotypically susceptible virions is $\gamma\beta\mu$. Since each of these virions represents an independent trial, the probability that all of them fail to establish a new infection (and subsequently result in the extinction of the population) is given by:

$$P(\text{res. extinct}) = (1 - y_0)^{\gamma\beta\mu} \approx e^{-y_0\gamma\beta\mu}.$$

Therefore, the threshold at which the extinction probability is 0.5 is given by $\gamma\mu = -\frac{\ln(0.5)}{y_0\beta}$. Using our empirically estimated values for $\beta$ and $y_0$, this predicts a 50% extinction probability when the product $\gamma \times \mu \approx 8.53$. Stochastic simulations approximately agree with this threshold, albeit with a slight upward shift, placing the inflection point at $\gamma \times \mu \approx 13$ (**Figure S9**). This modest discrepancy arises from two additional dynamics captured in the full simulation but not in this simplified analytical model. First, as cell entry is a binomial process, not every cell in the population will be infected, and therefore, the population size of replicating viruses will be diminished compared to our assumption. Second, resistant survival is not only contingent on surviving after initial emergence, as it may coinfect with susceptible genomes in subsequent cycles, potentially reducing the survival probability of its offspring.

#### Supplemental References


[1] Marc S. Collett et al. "Antiviral Activity of Pocapavir in a Randomized, Blinded, Placebo-Controlled Human Oral Poliovirus Vaccine Challenge Model". In: *The Journal of Infectious Diseases* 215.3 (Feb. 2017), pp. 335–343. ISSN: 0022-1899. DOI: 10.1093/infdis/jiw542.

[2] Michael B. Schulte and Raul Andino. "Single-Cell Analysis Uncovers Extensive Biological Noise in Poliovirus Replication". en. In: *Journal of Virology* 88.11 (June 2014). Ed. by S. Perlman, pp. 6205–6212. ISSN: 0022-538X, 1098-5514. DOI: 10.1128/JVI.03539-13.

[3] Olen M. Kew et al. "Vaccine-Derived Polioviruses and the Endgame Strategy for Global Polio Eradication". In: *Annual Review of Microbiology* (Oct. 2005). DOI: 10.1146/annurev.micro.58.030603.123625.

[4] Carlton E. Schwerdt and Jørgen Fogh. "The ratio of physical particles per infectious unit observed for poliomyelitis viruses". In: *Virology* 4.1 (Aug. 1957), pp. 41–52. ISSN: 0042-6822. DOI: 10.1016/0042-6822(57)90042-9.

[5] Andreas Dotzauer and Leena Kraemer. “Innate and adaptive immune responses against picornaviruses and their counteractions: An overview”. In: *World Journal of Virology* 1.3 (June 2012), pp. 91–107. ISSN: 2220-3249. DOI: 10.5501/wjv.v1.i3.91.

[6] Elizabeth J Tanner et al. “Dominant drug targets suppress the emergence of antiviral resistance”. In: *eLife* 3 (Nov. 2014). Ed. by Wenhui Li, e03830. ISSN: 2050-084X. DOI: 10.7554/eLife.03830.

[7] R Core Team. *R: A Language and Environment for Statistical Computing*. R Foundation for Statistical Computing. Vienna, Austria, 2024. URL: https://www.R-project.org/.

[8] Hadley Wickham. *ggplot2: Elegant Graphics for Data Analysis*. Springer-Verlag New York, 2016. ISBN: 978-3-319-24277-4. URL: https://ggplot2.tidyverse.org.

[9] Hong-Mei Liu et al. “Characterization of Poliovirus Variants Selected for Resistance to the Antiviral Compound V-073”. In: *Antimicrobial Agents and Chemotherapy* 56.11 (Nov. 2012), pp. 5568–5574. DOI: 10.1128/AAC.00539-12.

[10] Diana V Kouiavskaia et al. “Immunological and Pathogenic Properties of Poliovirus Variants Selected for Resistance to Antiviral Drug V-073”. en. In: *Antiviral Therapy* 16.7 (Oct. 2011), pp. 999–1004. ISSN: 1359-6535. DOI: 10.3851/IMP1838.

[11] Adi Stern et al. “The Evolutionary Pathway to Virulence of an RNA Virus”. English. In: *Cell* 169.1 (Mar. 2017), 35–46.e19. ISSN: 0092-8674, 1097-4172. DOI: 10.1016/j.cell.2017.03.013.

[12] Michael B Schulte et al. “Experimentally guided models reveal replication principles that shape the mutation distribution of RNA viruses”. In: *eLife* 4 (Jan. 2015). Ed. by Stephen P Goffs, e03753. ISSN: 2050-084X. DOI: 10.7554/eLife.03753.

[13] Nadège Néant et al. “Modeling SARS-CoV-2 viral kinetics and association with mortality in hospitalized patients from the French COVID cohort”. In: *Proceedings of the National Academy of Sciences* 118.8 (Feb. 2021), e2017962118. DOI: 10.1073/pnas.2017962118.

[14] Achim Zeileis and Gabor Grothendieck. “zoo: S3 Infrastructure for Regular and Irregular Time Series”. In: *Journal of Statistical Software* 14.6 (2005), pp. 1–27. DOI: 10.18637/jss.v014.i06.
